# Soybean PHR1-regulated low phosphorus-responsive *GmRALF22* increases uptake of phosphate via stimulating *GmPT*s expression

**DOI:** 10.1101/2024.05.29.596424

**Authors:** Fangjian Li, Cuishan Mai, Yan Liu, Yaru Deng, Lixia Wu, Xinni Zheng, Huijing He, Yilin Huang, Zhenxi Luo, Jinxiang Wang

**Affiliations:** Root Biology Center, South China Agricultural University, Guangzhou 510642, P.R. China; College of Natural Resources and Environment, South China Agricultural University, Guangzhou 510642, P. R. China; Key Laboratory of Agricultural and Rural pollution Control and Environmental Safety in Guangdong Province, Guangzhou 510642, P. R. China

**Keywords:** GmRALF22, GmPHR1, low phosphorus stress, GmPTs, soybean

## Abstract

Phosphorus (P) is one of essential macronutrients for plant growth and development. Rapid Alkalization Factors (RALFs) play crucial roles in plant responses to nutrient stresses, however, the functions of *Glycine max* RALFs (GmRALFs) under low P (LP) stress remain elusive. In this study, we first identified 27 *GmRALF*s in soybean, then we revealed that *GmRALF10, GmRALF11*, and *GmRALF22* are induced in both roots and leaves, while only *GmRALF5*, *GmRALF6*, and *GmRALF25* are up-regulated in leaves in LP conditions. Furthermore, *GmRALF22* was found to be the target gene of transcription factor *GmPHR1,* which binds the P1BS *cis*-element in the promoter of *GmRALF22* via electrophoretic mobility shift assay (EMSA) and DUAL-LUC experiment. Colonization of *Bacillus subtilis* that deliver GmRALF22 increases the expression of high affinity phosphate (Pi) transporter gene *GmPT2, GmPT11, GmPT13* and *GmPT14*, thus increases the total amount of dry matter and soluble Pi in soybean. RNA-sequencing uncovered that GmRALF22 alleviates LP stress by regulating the expression of JA-, SA- and immune-related genes. Finally, we verified that GmRALF22 is dependent on FERONIA to promote Arabidopsis primary root growth under LP conditions. In summary, *GmPHR1-GmRALF22* module positively regulates soybean’s tolerance to LP.

**Highlights:**

1. Soybean genome has 27 *GmRALF*s. *GmRALF5*, *GmRALF6, GmRALF10*, *GmRALF11*, *GmRALF22* and *GmRALF25* are induced in low phosphorus (LP).
2. GmPHR1 directly regulate the transcription of *GmRALF22 via* binding the promoter P1BS *cis*-element.
3. Secretion of GmRALF22 protein by *Bacillus subtilis* promoted soybean growth under LP conditions by improving soybean P nutrition through increased expression of high affinity phosphate (Pi) transporter gene.
4. GmRALF22 regulates soybean P nutrition by JA-, SA- and immune-related genes expression at transcriptome level. That application of GmRALF22 promoted primary root growth in Arabidopsis in LP is dependent on FER receptor.

## 1. Introduction

Phosphorus (P) is an indispensable nutrient element for plant growth and development, hence, the deficiency of P in soils results in the decrease of crop yield and quality (Ding et al., 2021; Dissanayaka et al., 2018; Hu et al., 2011; Liu et al., 2018 Prathap et al., 2022). Soybean (*Glycine max L*) is one of important grain and oil crops with remarkable nutritional value worldwide (Cao et al., 2021; Jian, 2020). As biological nitrogen fixation and phospholipid biosynthesis consume phosphate (Pi), soybean is prone to be adversely affected by insufficient P availability in nature. Therefore, it is imperative to investigate the underlying mechanisms governing soybean’s response to low P (LP) stress.

P deficiency can result in nutrient imbalance within plants, characterized by an augmented abundance and density of lateral roots and root hairs, darkened leaf pigmentation, delayed growth, and flowering, as well as reduced plant stature (Pérez-Torres et al., 2008). Concurrently, a cascade of physiological and biochemical responses is triggered in plants including phospholipid degradation, enhanced synthesis of glycolipids and sulfur lipids, ultimately leading to diminished crop yield (Prathap et al., 2022; Wang et al., 2023).

Phosphate Starvation Response 1 (PHR1) is a major transcription factor that responds to LP stress and regulates the P-related signaling (Sega et al., 2019). Under LP conditions, PHR1 binds to the *cis*-element P1BS in the promoter of *P Starvation-Induced* (*PSI*) to activate downstream gene expression (Xue et al., 2017). The genes activated by PHR1 can systematically or locally participate in regulating P absorption and assimilation, as well as participating in synthesizing enzymes involved in nucleic acid synthesis, galactose, and redistributing P by affecting P transporters (Zhou et al., 2021). The Arabidopsis genome also has PHR1-like genes such as *PHL1, PHL2* and *PHL3* (Wang et al., 2023). Of note, a total of 35 homologous genes of *PHR1* exists in soybean genome (Xue et al., 2017). Moreover, transcriptome analysis of sorghum *PHR1* transgenic material and *phl1* mutant growing under P starvation shows that *PHR1* also affects the expression of genes related to metabolic pathways of ethylene, jasmonic acid (JA), and salicylic acid (SA), which interact with each other and play important roles in plant stress resistance (Zhang et al., 2019).

As crucial signaling molecules, signal peptides interact with their corresponding receptors or co-receptors to establish intricate networks that exert a profound influence on plant growth, development, and adaptive responses to dynamic environments (Murphy et al., 2014; Ogilvie et al., 2014). The identified signal peptides can be primarily categorized into two groups: small post-translationally modified peptides and cysteine-rich polypeptides (Murphy et al., 2016). Rapid alkalization factor (RALF) belongs to the cysteine-rich polypeptide family in plants. It was initially identified for its ability to increase pH levels in tobacco suspension cell culture media (Pearce et al., 2001). The N-terminal of mature RALF typically contains conserved YISY domains required for binding receptors, which are essential for RALF’s function. Additionally, there is a dibasic site called "RR", which is significant for processing RALF into mature peptide (Pearce et al., 2010). For instance, the "RR" dibasic site upstream of AtRALF1 is so critical that mutation leads to decreased accumulation of mature AtRALF1 (Matos et al., 2008). The investigation of RALF genes has been carried out in diverse plant species, including *Arabidopsis thaliana*, tobacco, and tomato, encompassing various physiological and developmental processes such as root development (Wu et al., 2007), sexual reproduction (Covey et al., 2010), acidification (Yang et al.,2022), cell expansion and immune responses (Tang et al., 2022). A lot of studies have revealed that RALF is involved in various aspects of plant growth and development by reacting/binding/acting with FERONIA (FER) kinase, the receptor of RALF (Tang et al., 2022). AtRALF1 and AtFER promote the growth of root hair tip by regulating protein synthesis (Zhu et al., 2020). In addition, the expression of Arabidopsis *RALF* is induced by low P conditions, which triggers the immune responses by PHR1-RALF-FERONIA pathway and changes the microbial environment in the rhizosphere in *Arabidopsis thaliana*, thus alleviating P starvation (Tang et al., 2022). The overexpression of *RALF1* in *Arabidopsis thaliana* impedes cellular expansion and root elongation, potentially influenced by the brassinolide-mediated signaling pathway (Zhao et al., 2021). As reported, a total of 18 GmRALFs respond to Fusarium oxysporum (F. oxysporum) infection (Liu et al., 2022), indicating the roles of GmRALF in immunity. However, the function of RALF in soybean remains elusive.

Previous studies have demonstrated the pivotal role of RALFs in mediating responses to nutrient stresses (Fu et al., 2020; Wang et al., 2018), however, investigations specifically focusing on soybean RALF remain limited. In this study, we firstly identified soybean RALF genes at genome level through bioinformatics analysis. Then we found that *GmRALF10*, *GmRALF11*, *GmRALF22* are induced by LP in both roots and leaves, and *GmRALF5*, *GmRALF6*, and *GmRALF25* are only induced in leaves via qRT-PCR. Then DUAL-LUC and EMSA experiments verified that GmPHR1 positively regulates the expression of *GmRALF22* through directly binding to the promoter P1BS *cis*-element. Furthermore, the inoculation of *B. subtilis* that secrets GmRALF22 protein enhances soybean growth under LP conditions by promoting uptake of P through up-regulating the expression of high-affinity phosphate (Pi) transporter genes *GmPT2*, *GmPT7*, *GmPT11*, and *GmPT14*. RNA-seq analysis suggests that GmRALF22 regulate soybean P nutrition at the transcriptional level via modulating JA-, SA-, and immune-related gene expression. In addition, GmRALF22 facilitates primary root growth in *Arabidopsis thaliana* in LP is dependent on the FERONIA (FER) receptor protein.

## 2. Materials and methods

### 2.1 Bioinformatics analysis of GmRALF family members

The protein sequences of Arabidopsis RALFs (AtRALF1*∼*AtRALF34) were obtained from TAIR database (https://www.arabidopsis.org/). After blast against soybean proteome (Wm82.a4. v1) with AtRALFs, the sequences of GmRALF family members were retrieved from Phytozome (https://phytozome-next.jgi.doe.gov/). The ExPasy tool (https://web.expasy.org/protparam/) was utilized to predict the physical and chemical characteristics of the *GmRALF* family, including molecular weight, theoretical isoelectric point, hydrophilicity coefficient, and instability coefficient. The subcellular localization of the GmRALF family members were predicted using Cell-PLoc 2.0 (http://www.csbio.sjtu.edu.cn/bioinf/plant-multi/). *GmRALF*s were named based on their physical location on soybean chromosome.

### 2.2 Phylogenetic tree reconstruction of GmRALFs in soybean

The amino acid sequences of GmRALFs were obtained from soybean (Wm82.a4. v1) and *Arabidopsis thaliana* (TAIR 10) proteosome from Phytozome (https://phytozome-next.jgi.doe.gov/). The phylogenetic tree was reconstructed by MEGA 7.0 program using the neighbor-joining method, with 1000 bootstrap replicates. The bootstrap values are indicated for major branches as percentages.

The conserved motifs of GmRALFs were analyzed with MEME suite (suite.org/meme/tools/meme). The Batch CD Search tools from the NCBI website (https://www.ncbi.nlm.nih.gov/Structure/bwrpsb/bwrpsb.cgi) were used to analysis the conserved domain of GmRALF family.

### 2.3 Analysis of P1BS cis-elements in GmRALF promoter

The promoter sequences (2000 bp upstream of the initiation codon) of each member of the GmRALF family in soybean genome (Wm82.a4. v1) are downloaded) from Phytozome website (https://phytozome-next.jgi.doe.gov/). Then the P1BS *cis*-element in the promoter region of *GmRALF*s were screened at PlantCARE website (http://bioinformatics.psb.ugent.be/webtools/plantcare/html/)

### 2.4 Plant materials and growth

The *Glycine max* genotype Yuechun 03-3 (YC03-3) was used. Soybean seeds were sterilized with chlorine (100 mL NaClO + 4.2 mL 36%∼38% HCl) for 4 hours. The sterilized seeds were evenly planted in clean and moist quartz sand, and grown in growth room (25 ℃, light 16 h/ dark for 8 h) for 4 days. The uniform seedlings were transplanted into nutrient solution containing 500 μM phosphate (HP) or 5 μM phosphate (LP) (Ou et al., 2022). The nutrient solution was changed every seven days. After 10 days of treatment, appropriate amount of roots and leaves were put into liquid nitrogen, and quickly stored at -80 ℃.

The *Arabidopsis thaliana* (L.) wild type Columbia (Col-0) was used. The *fer4* mutant (a loss-of-function mutant of *FER*) seeds were kindly provided by Dr. Fen Yu (Tang et al, 2022). Arabidopsis seeds were sterilized with 75% alcohol for 3 minutes, soaked in 95% alcohol for 30 seconds, and dried. They were then sowed on half strength Murashige and Skoog (1/2 MS) media containing 0.8% sucrose and 1% Phytagel (Sigma-Aldrich), followed by a stratification treatment at 4 ℃ in the dark for two days. Finally, they were placed in a growth chamber with a thermocycle of 22 ℃/20 ℃, a photoperiod of 16 h light/8 h dark, and a light intensity of 100 μmol^-2^.

### 2.5 RNA extraction, reverse transcription, and qRT-PCR analysis

Total RNA was extracted from leaves and roots of soybean by Trozol. Total RNA Extraction Reagent (Genesand, China, RE703 https://www.genesand.com/). RNA was reverse-transcribed into cDNA by using reverse transcription kit (ABclonal, China, RK20429 https://abclonal.com.cn/).

After total RNA extraction and cDNA systhesis were carried out according to the method as described (Li et al., 2023). The cDNA was used for quantitative qRT-PCR analysis of SYBR Green monitoring on the 7500 real-time PCR system (Thermo, USA). *Glycine max EF1a* (*GmEF1a, Glyma.17G186600*) was used to normalize qRT-PCR data (Xu et al., 2013). All specific qRT-PCR primer pairs are listed in Table S1.

### 2.6 Vector of construction

Using soybean cDNA template, the cDNA sequence (open reading frame) of *GmPHR1* (*Glyma.01G009600*) was amplified with specific primer pair, then *GmPHR1* was cloned into the expression vector pCambia1300 digested with SalI and PstI, respectively. In addition, the cDNA of *GmPHR1* was cloned into the prokaryotic expression vector pGEX-6p-3(GE Healthcare, USA) cleaved with BamHI. The promoter sequence of *GmRALF22* was amplified with specific primers o using soybean genomic DNA as template, thus the promoter sequence of *GmRALF22* was cloned into the LUC tag-fused vector pGreen 0800 II. All specific clone primer pairs are listed in Table S1.

### 2.7 Application of GmRALF22

The mature GmRALF22 protein was synthesized by Sangon Biotech (China, www.sangon.com). The protein powder was dissolved in sterilized water to the concentrations as indicated. As to RALE-mediated root growth assay, Arabidopsis seeds were germinated on 1/2 MS agar plates for three days before infiltrating each plant’s roots with 50 μl of a 2 μM GmRALF22 protein solution. After five days of growth, the number of lateral roots in *Arabidopsis thaliana* was quantified using a stereoscopic microscope (Evidengt, Germany). After capturing images of the samples using a Nikon D300s digital camera, the length of the primary root was quantified with ImageJ software (Ou *et al*.,2022).

### 2.8 B. subtilis-GmRALF22 treatment

HP and LP Hoagland nutrient solutions were mixed with fertilizer-free substrate (Jiffy, Estonia) to form HP and LP growth media, respectively. The height of the pots for culturing soybean seedlings is 13 cm and the diameter is 20 cm. Based on native GmRALF sequences, we optimized *GmRALF* codons (Seqid No.1) to increase the translational efficiency of *GmRALF* in *B.subtilis* and cloned it into the plasmid pBEs, which was introduced into *B. subtilis* RIK1285 (Tang et al.,2022). The transformed *B. subtilis* RIK1285 were cultured until the OD600 reach to 0.6, thus the bacteria were inoculated into soybean roots grown in LP for one week, and the empty vector *B. subtilis* was set up as control.

### 2.9 Dual-luciferase (DUAL-LUC) assay

*Nicotiana benthamiana* seedlings were cultivated in soils for 4-5 weeks. The Agrobacterium GV3101 strain containing the ProGmRALF22-LUC or GmPHR1 constructs was inoculated in YEP medium and incubated at 28°C for 24 hours. Transient expression mediated by *Agrobacterium tumefaciens* was performed in *Nicotiana benthamiana* leaves following the protocol described by (Li et al., 2023). After two days, fluorescence in transformed *Nicotiana benthamiana* was detected using a *in vivo* imager.

The LUC and REN activities was measured with Dual Luciferase Reporter Gene Assay Kit (YeSen, China https://www.yeasen.com/) as described (Li et al., 2023). The 100 mg of fresh leaf powder was suspended in 100 μL of lysis buffer on ice for 5 minutes and then centrifuged at a speed of 13,000 r/min for 1 minute. A volume of 20 μL of the supernatant was transferred into a mixture containing 100 μL of firefly luciferase detection reagent to measure LUC activity using a multifunctional microplate reader (Turner Biossystem). Subsequently, the above mixture was combined with 100 μL of REN test working solution to determine REN activity. The relative activity ratio between LUC and REN was calculated as LUC/REN. Three biological repeat sequences were measured in each sample and similar results were obtained.

### 2.10 Determination of height, fresh weight, dry weight, and soluble phosphate concentration

The height of soybean seedlings was measured on the 11^th^, 14^th^,17^th^, 21^th^ day (after application of *B. subtilis*) under HP or LP condition. The fresh weight (FW) of soybean seedlings was weighted on the 30^th^ day under HP or LP conditions. The dry weight (DW) of soybean seedlings was measured after drying at 105°C for 30 min, and then oven-drying at 75°C. To measure soluble Pi concentration, about 0.1 g of fresh sample was ground in deionized water, and the supernatant was used for determining the amount of extracted Pi as described before (Murphy and Riley, 1962)

### 2.11 Electrophoretic mobility shift assay (EMSA)

EMSA was carried out according to the manufacturer’s protocol using the EMSA/Gel- Shift kit (GS009, Beyotime, https://www.beyotime.com/index.htm, China) as described (Li et al., 2023). GST-GmPHR1 protein and GST protein were used for EMSA assay. The primers of *GmRALF22* were synthesized and labelled with biotin by Sangon Biotech (China, www.sangon.com). All EMSA primer pairs are listed in Table S1.

### 2.12 RNA sequencing analysis of soybean roots inoculated with B. subtilis that secrets GmRALF22

Approximately 0.5 g roots were weighted from soybeans treated with *B. subtilis* transformed with empty vector (*B. subtilis* EV) or expressing GmRALF22 vector (*B. subtilis* GmRALF22) under LP conditions, with three replicates per treatment. A total of six samples were sent to Gene denovo (https://www.genedenovo.com/) for sequencing library construction and RNA-Seq analysis. Differentially expressed gene (DEG) were determined according to p < 0.05, fold changes >2. Gene Ontology (GO), and Kyoto Encyclopedia of Genes and Genomes (KEGG) were analysis as described (Xie et al., 2022).

### 2.13 Protein extraction and immunoblot analysis

Total protein was extracted as described before (Tang et al., 2022). *B. subtilis*- GmRALF22 was cultured in LB culture at 37℃ and 200 r/min for 8 hours. The bacteria were collected by centrifugation and diluted with water at 10: 1.and mixed liquor was loaded on 10% SDS-PAGE for Western Blotting using anti-His (M30111) (Abmart, http://www.ab-mart.com.cn/, China).

### 2.14 Data processing and analysis

All data were analyzed with Microsoft Excel 2016 (Microsoft Company, USA) software. Student’s *t*-test was employe to compare differences. All charts were drawn with Microsoft Excel 2016 or OriginPro 2022.

## 3 Results

### 3.1 Identification and phylogenetic analyses of Glycine max RALF members

As shown in Table S2, a total of 27 *GmRALFs* has been found in soybean genome. *GmRALF*s are distributed on chromosome 1, 3, 5, 7, 8, 10-13, and 15∼20, respectively. Accordingly, they were named as *GmRALF1-GmRALF27,* respectively (Table S2). The number of amino acids of GmRALFs ranges from 73 to 174, with a molecular weight spanning from 6.04 to 19.73 KDa (Table S2). The theoretical isoelectric points of all members of GmRALF family were greater than 7, which indicating that GmRALFs were basic proteins (Table S2). The hydrophilic coefficient of GmRALF family members ranges from -0.981 to 0.079, and the instability coefficient of GmRALF members are from 19.69 to 73.83 (Table S2).

The subcellular localization of GmRALFs were predicted with Cell-PLoc 2.0. As shown, GmRALF9, GmRALF21, GmRALF22, GmRALF25 appear to be located on membrane or nucleus; GmRALF4 and GmRALF16 may be located on membrane, chloroplast or nucleus; GmRALF10 is located on cell wall, chloroplast, nucleus; GmRALF12 seem to be located on chloroplast, mitochondria or nucleus; GmRALF26 is located on chloroplast, nucleus;GmRALF13, GmRALF24, GmRALF27 may be located on membrane, the others may be only located on nucleus (Table S2).

To analyzes the evolutionary relationships of different RALF proteins in soybeans, RALFs in *Arabidopsis* and soybean were selected, the phylogenetic trees were reconstructed by neighbor-joining method. Soybean RALF are mainly divided into three subfamilies. Among them, *GmRALF7, GmRALF11*, *GmRALF12*, and *GmRALF21* belong to the first subgroup; *GmRALF17*, *GmRALF26* are grouped into second subgroup; *GmRALF1, GmRALF2, GmRALF3, GmRALF4, GmRALF5, GmRALF6, GmRALF8, GmRALF9, GmRALF10, GmRALF13, GmRALF14, GmRALF15, GmRALF16, GmRALF18, GmRALF19, GmRALF20, GmRALF22, GmRALF23, GmRALF24, GmRALF25, GmRALF27* are the members of the third subgroup (Fig. 1). The high similarity of RALF family between soybean and *Arabidopsis* suggests that RALF protein is conserved in higher plants.

**Fig. 1.**
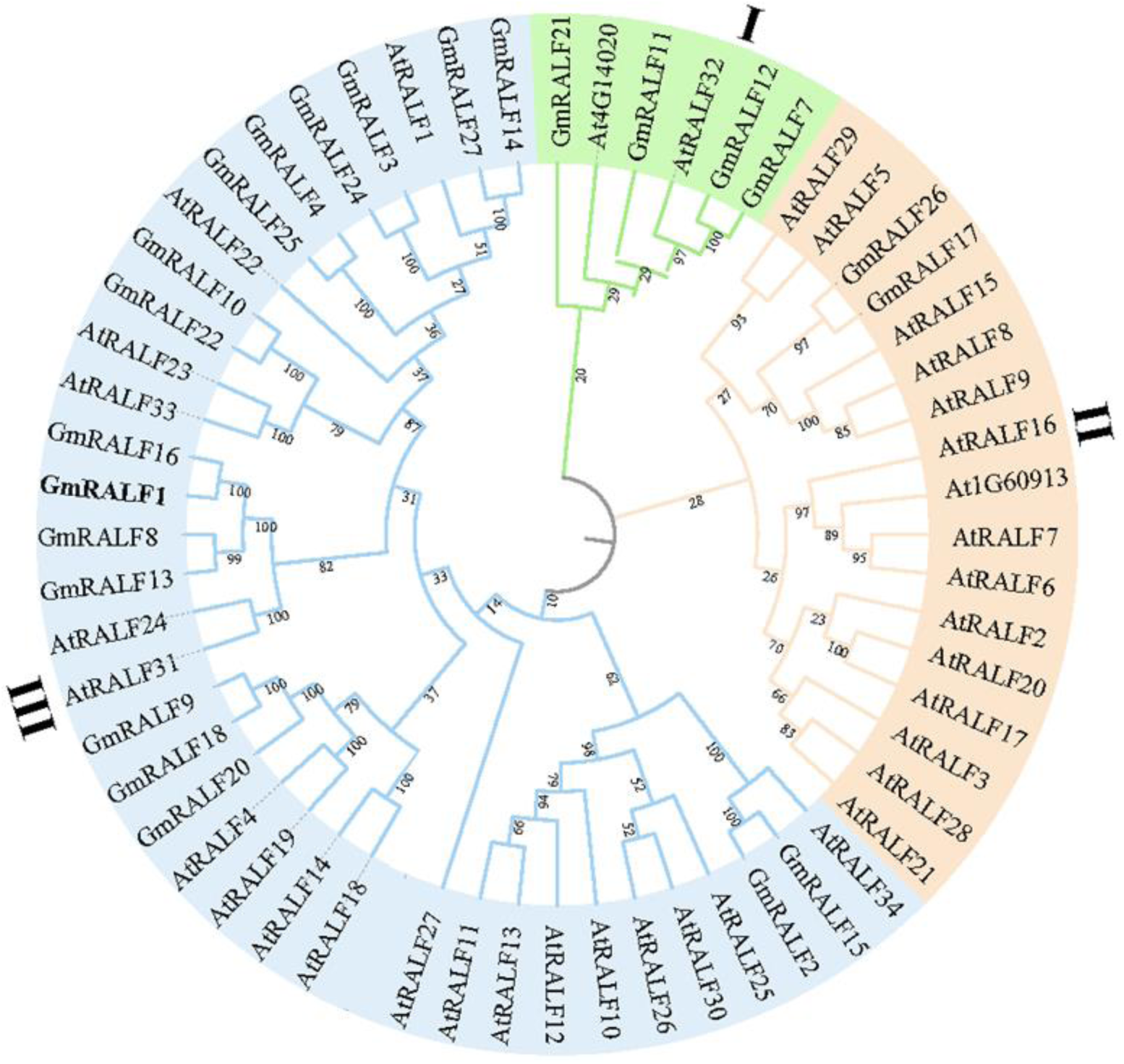
Phylogenetic analysis of RALF proteins from *Glycine max* (*Gm*) and *Arabidopsis thaliana* (*At*). The phylogenetic tree was reconstructed with MEGA 7.0 based on the neighborhood method. The bootstrap value was set to 1000.

To further understand the function of GmRALFs, conserved motifs and domains were analyzed. The results shows that a total of 18 GmRALFs contain three conserved motifs, and 20 GmRALFs contain two conserved motifs and 7 GmRALFs only contain motif 1 (Fig. S1A). Of important, Motif 1 contains the “YISY” sequence, which is necessary for RALF to bind to receptors (Pearce et al., 2010). Motif 3 contains the two base sites required for RALF to be processed into mature peptides, named “RR” (Pearce et al., 2010). Of note, a total of 20 GmRALFs (GmRALF1, GmRALF4*-*GmRALF5, GmRALF6*-*GmRALF9, GmRALF11- GmRALF17, GmRALF19, GmRALF21-GmRALF25, GmRALF27) contain motif 1, respectively. In addition, a total of 22 GmRALFs such as GmRALF1, GmRALF4*-* GmRALF15, GmRALF17, GmRALF18*-*GmRALF25, and GmRALF27 have “RR”, (Fig. 2).

**Fig. 2.**
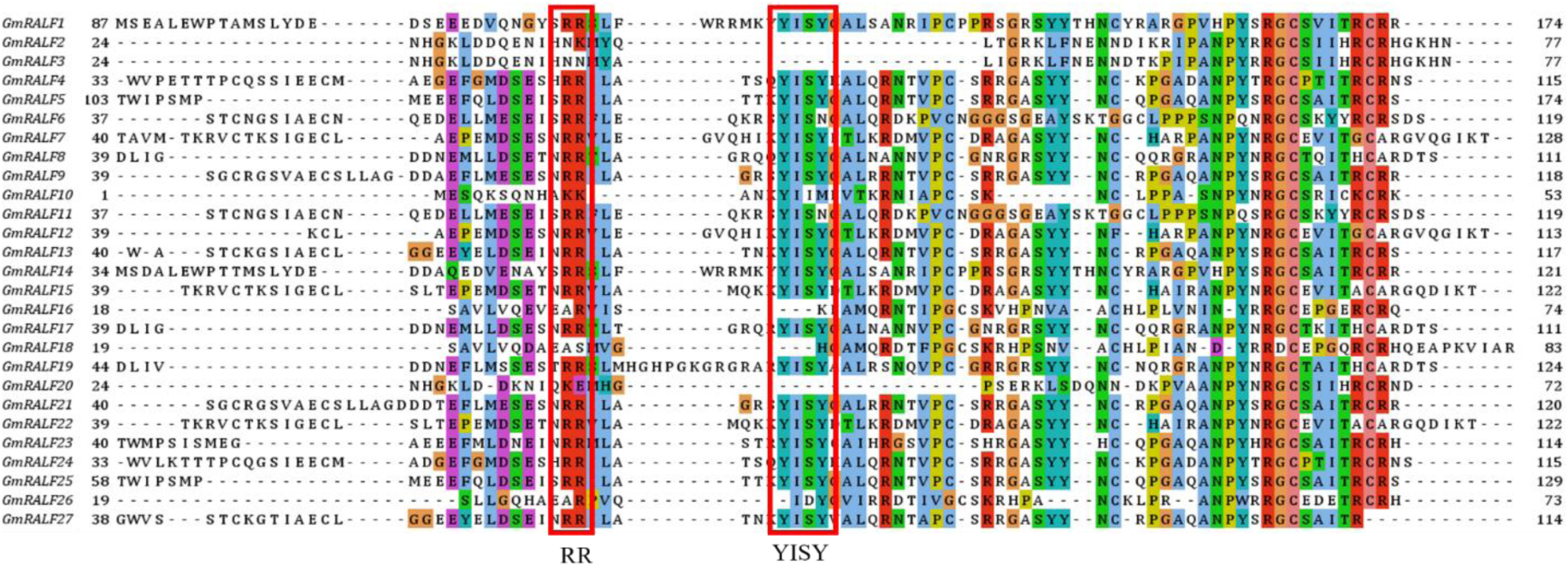
The alignment of *Glycine max* RALFs amino acid sequence. The clustalw programme was used. The conserved motifs “RR” and YISY were highlighted with red square.

### 3.2 Expression pattern of GmRALFs

By querying the database of Soybase (www.soybase.org), we found the expression levels of 19 *GmRALF*s in different tissues of soybean, and constructed the heat map based on these data (Fig. 3A). Among them, the relative expression level of *GmRALF14* is high in all soybean tissues. That of *GmRALF1* is high in other tissues except root nodules, and *GmRALF27* has a high expression level in other tissues except 42 DAF seeds (Day after flowering) seeds (Fig. 3A). The expression levels of *GmRALF4*, *GmRALF6*, *GmRALF7* and *GmRALF24* are relatively low in tested soybean tissues; the expression levels of *GmRALF8*, *GmRALF16*, *GmRALF17* and *GmRALF19* are low in other parts of soybean except flowers, and the expression levels of *GmRALF10* are low in other parts of soybean tissues except roots; the expression level of *GmRALF9* in 10-21 DAF seeds are mild, but higher in other parts; *GmRALF21* only had a high expression level in 14 DAF pods. *GmRALF5*, *GmRALF15* and *GmRALF22* are expressed at a moderate level in soybean tissues (Fig. 3A).

**Fig. 3.**
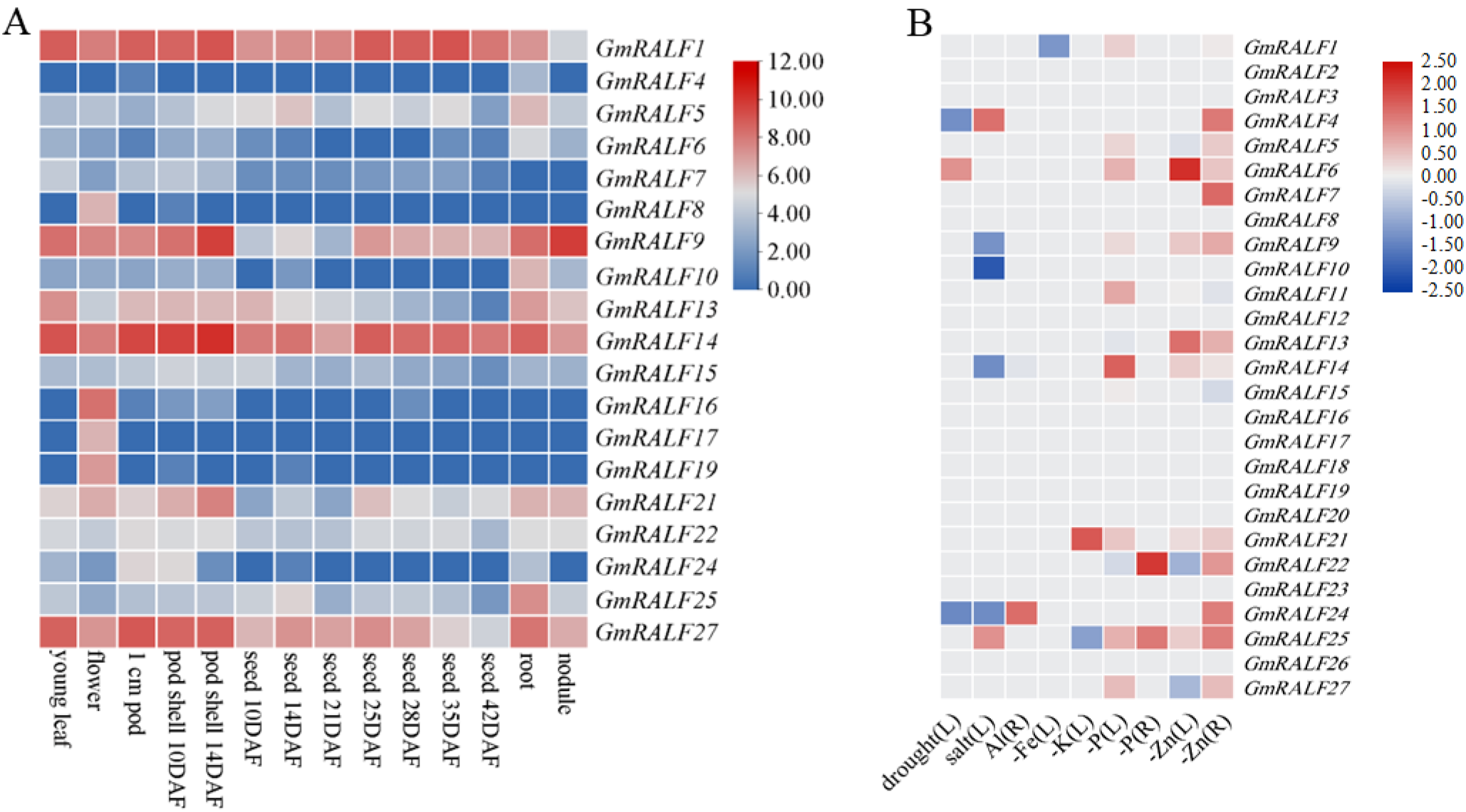
Expression patterns of *GmRALFs* in different soybean organ (A) and different abiotic stresses (B). The relative expression levels of *GmRALFs* were retrieved from published RNA-seq data (Belamkar et al, 2014; Zhao et al, 2020; Kohlhase et al, 2021; Wang et al, 2012; Mo et al, 2019; Zeng et al, 2018; Zeng et al, 2019), and the heatmap was drawn with TBtools (Chen et al.,2023). (A) The relative expression levels of *GmRALFs* gene in different organ of soybean; (B) The relative expression level of *GmRALFs* gene under different abotic stress. The color scale on the right of Figure 3A and Figure 3B indicates the relative expression level (log2(RPKM)) of *GmRALFs*. RPKM: Reads per kilobase/million.

We date-mined and obtained the expression levels of *GmRALF* genes under drought, salt (Belamkar et al, 2014), Al toxicity (Zhao et al, 2020), Fe deficiency (Kohlhase et al, 2021), potassium deficiency (K) (Wang et al, 2012), P deficiency (Mo et al, 2019; Zeng et al, 2018) and Zn deficiency (Zeng et al, 2019). Under drought conditions, the expression levels of *GmRALF4* and *GmRALF24* are down-regulated and that of *GmRALF6* is up-regulated in soybean leaves. In salt-stressed leaves, the transcripts of *GmRALF4* and *GmRALF25* is up- regulated, while that of *GmRALF9*, *GmRALF10*, *GmRALF14* and *GmRALF24* is down- regulated; *GmRALF24* is up-regulated in roots under Al treatment. *GmRALF1* is induced by low Fe in leaves. *GmRALF21* is up-regulated in K-deficient leaves, while *GmRALF25* is down-regulated. The expression of *GmRALF1*, *GmRALF5*, *GmRALF6*, *GmRALF9*, *GmRALF11*, *GmRALF14*, *GmRALF21*, *GmRALF25* and *GmRALF27* is up-regulated in low P stressed leaves. Similarly, the expression of both *GmRALF22* and *GmRALF25* were up- regulated in roots under P deficient conditions. Furthermore, *GmRALF6*, *GmRALF9*, *GmRALF13*, *GmRALF14*, *GmRALF21* and *GmRALF25* are found to be up-regulated in Zn-deficient leaves; whereas *GmRALF22*, *GmRALF5* and *GmRALF27* are down regulated. *GmRALF4, GmRALF9, GmRALF13, GmRALF14, GmRALF21, GmRALF22, GmRALF24, GmRALF25,* and *GmRALF27* are all observed to be up regulated in roots under Zn deficiency treatment (Fig.3B). These results indicate that *GmRALF*s play roles in soybean to cope with drought, salt, and nutrient deficiency.

### 3.3 Responses of GmRALFs to low P

P1BS (GNATATNC) element is the binding site of PHR1, which is a key transcription factor in P signal transduction (Wang et al., 2021). There are P1BS elements in the promoter regions of phosphate transporters induced by low P in different species (Karthikeyan *et al*., 2009; Wang et al., 2021). As revealed, GmPHR1 also binds to P1BS (He *et al*., 2023). Thus, the promoter region of *GmRALF* genes, encompassing approximately 2000 bp upstream from the ATG start codon, was retrieved from Phytozome and subjected to analyze the presence of P1BS binding sites associated with low P stress. As shown in Fig. 4A, the P1BS binding sites in the promoter region of *GmRALF5*, *GmRALF6*, *GmRALF10*, *GmRALF11*, *GmRALF20*, *GmRALF22*, *GmRALF25* were found (Fig. 4A). Of course, we can not rule out that other *GmRALF*s that lack the P1BS *cis*-element also respond to low P stress.

**Fig. 4.**
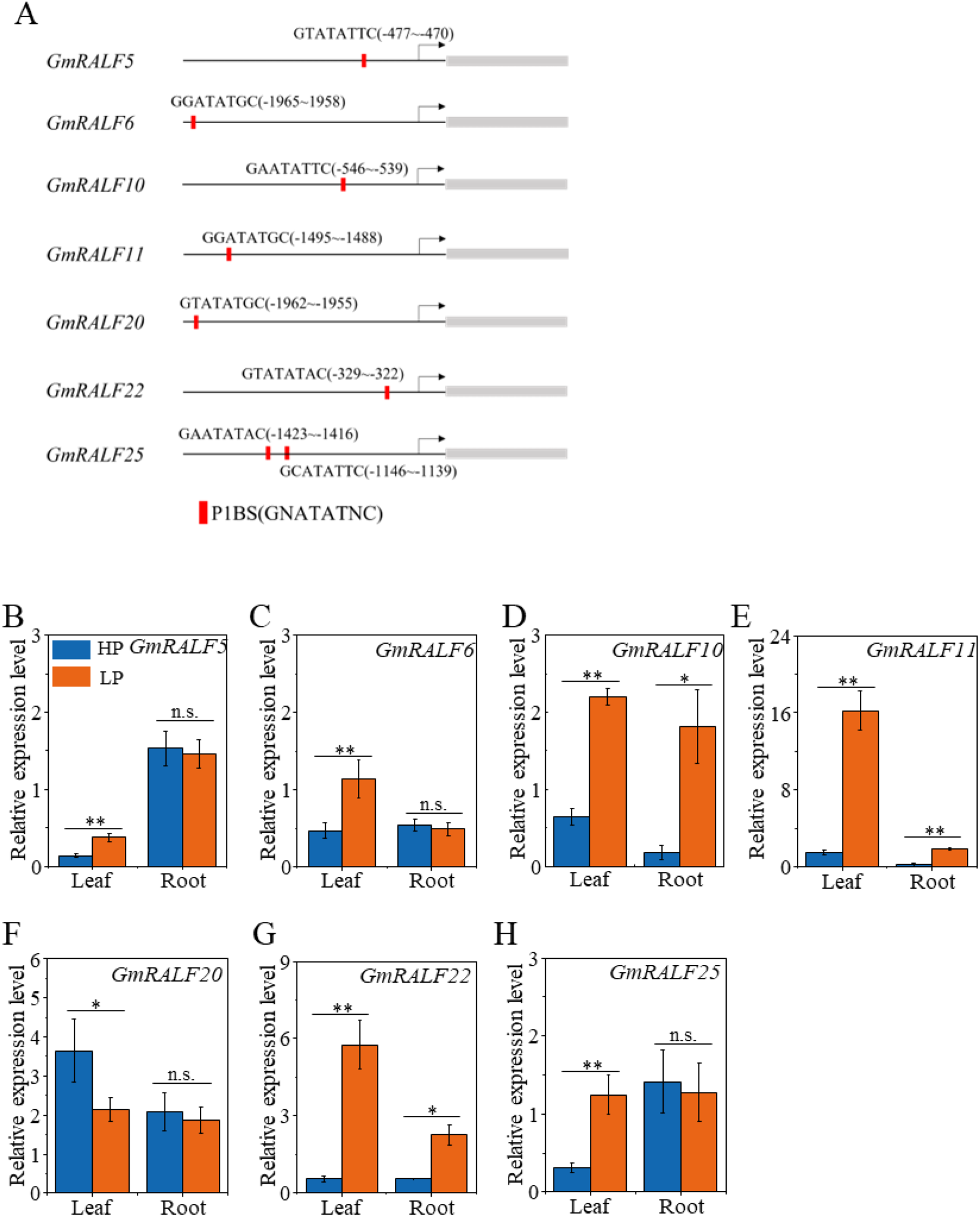
Analysis of P1BS *cis*-element in the promoter of *GmRALFs* and expression level of *GmRALFs*. (A) The predicted GmPHR1-binding sites (P1BS, GNATATNC) within 2,000 bp of *GmRALF* promoters; (B-H) The responses of *GmRALF5* (B), *GmRALF6* (C), *GmRALF10* (D), *GmRALF11* (E), *GmRALF22* (F), *GmRALF25* (G) to low P (LP) stress. The relative expression level of *GmRALFs* were normalized against *GmEF1a* (*Glyma.17G186600*). The high phosphorus (HP) and LP indicates the 500 μM and 5 μM phosphate, respectively. Data are means ± SE (n = 4). Student’s *t*-test was used to compare the difference between HP and LP (*, P < 0.05; **, P < 0.01; n.s., no significance).

To validate their responses to low P, the *GmRALF*s containing P1BS elements were subjected to qRT-PCR analysis. As shown, *GmRALF5*, *GmRALF6* and *GmRALF25* are up- regulated in leaves by 1.70, 1.45, 3.04 times by low, P respectively (Fig. 4B, C, H), while *GmRALF10*, *GmRALF11* and *GmRALF22* are all up-regulated by low P in leaves and roots, and *GmRALF10* is up-regulated by 2.40 times in leaves and 8.79 times in roots (Fig. 4D, E, G). *GmRALF11* is up-regulated by 10.23 times in leaves and 5.96 times in roots (Fig. 4E).

*GmRALF22* is up-regulated by 9.89 times in leaves and 3.07 times in roots in low P (Fig. 4G).

### 3.4 Transcript factor GmPHR1 activates GmRLAF22 transcription

Given that soybean GmPHR1 (Glyma.19G167500) is a MYB transcription factor, which is up-regulated under low P conditions (Li et al., 2022), and Arabidopsis *AtRALF1* is regulated by AtPHR1 (Tang et al., 2022). We thus hypothesized that soybean GmPHR1 may directly modulate the transcription of *GmRALF22*. To further investigate the involvement of GmPHR1 in the response of *GmRALF22* to low P mediated by P1BS elements, the effector containing an *GmPHR1* driven by constitutive 35 promoter and the reporter that contains LUC reporter gene driven by GmRALF22 promoter was constructed and introduced into GV3101, respectively (Fig. 5A).After co-transformation into tobacco leaves, the results demonstrated that the ectopic expression of *GmPHR1* significantly enhanced fluorescence intensity (Fig. 5B), the LUC/REN of the experimental group was 1.62 times higher (P < 0.05) than that of the control (Fig. 5C). This verify that GmPHR1 activates the activity of *GmRALF22* promoter *in vivo*.

**Fig. 5.**
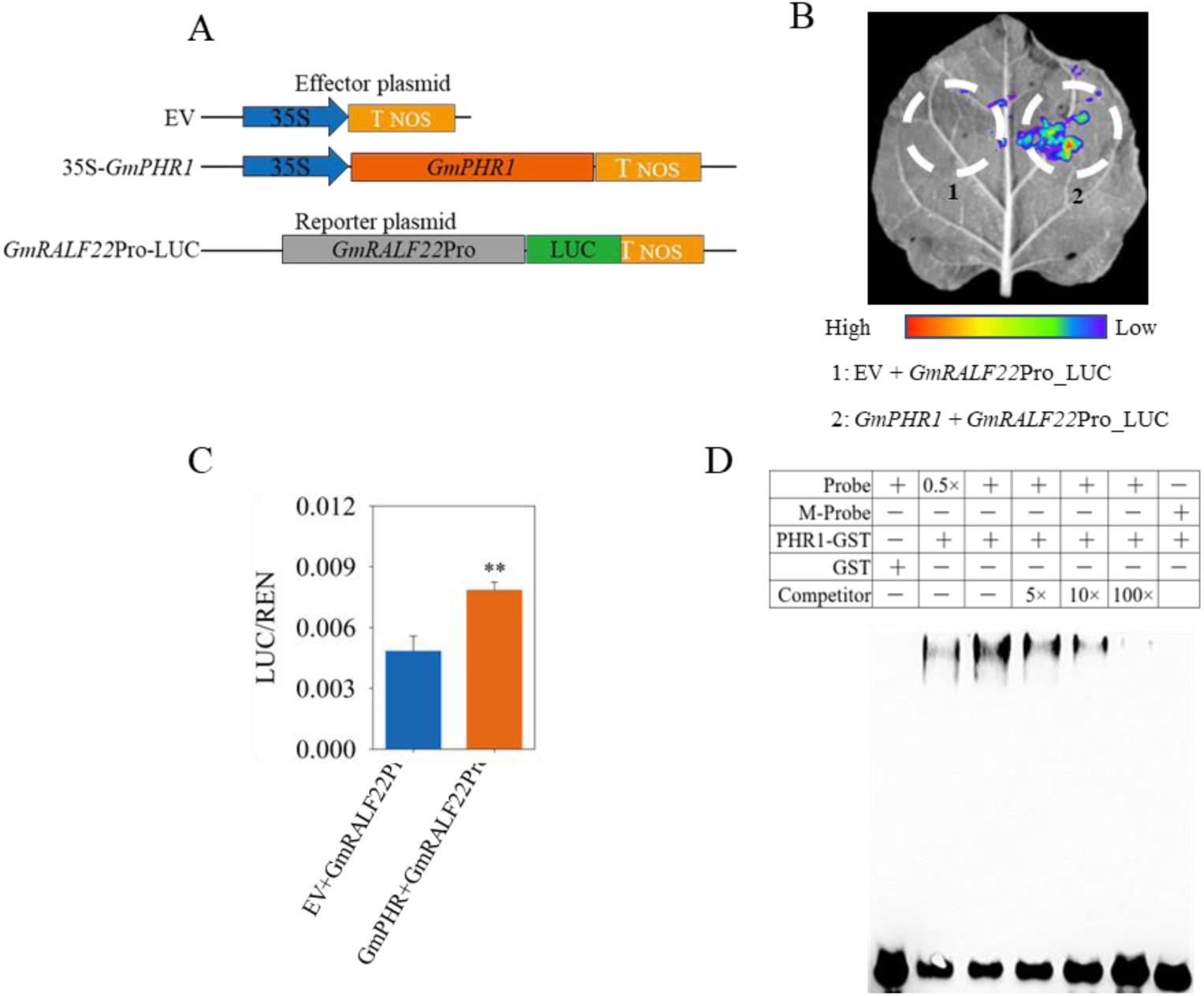
GmPHR1 promotes the expression of *GmRALF22* by directly binding its promoter *in vivo*. (A) Schematics of all constructs used for transient expression assays in *Nicotiana benthamiana* leaves. The *GmRALF22* promoter was fused to the luciferase (LUC) reporter gene. 35S: *GmPHR1* acted as an effector, and the empty vector (EV) acted as the negative control. (B) LUC image verify GmPHR1 activates *GmRALF22* promoter (Pro) activities. (C) The ratio of LUC to REN was quantified. (D) Electrophoretic mobility shift assay (EMSA) showed that GmPHR1 protein binds to the P1BS (GTAGAGAC) element in biotin-labelled *GmRALF22* probe. Data are means ± SE (n = 4). Student’s *t*-test was used to compare the difference between HP and LP (**, P < 0.01; n.s., no significance).

In addition, we verified the binding of GmPHR1 with *GmRALF22* probe *in vitro*. The purified GST-GmPHR1 protein and biotin-labeled *GmRALF22* probe (including P1BS element "CTATATAC") were used for EMSA. The results showed that GmPHR1 binds to the biotin-labeled probe containing P1BS elements, but not to the P1BS mutation probe ("TCCGCGGA"), and the addition of 100-fold cold probe significantly inhibited the binding of biotin-labeled probe to GmPHR1 (Fig. 5D). Collectively, these results show that GmPHR1 directly binds the P1BS element of *GmRALF22* promoter and positively promotes the expression of *GmRALF22 in vivo* and *in vitro*.

### 3.5 Bacterial colonization and delivery of GmRALF22 by B. subtilis alleviates LP stress

To explores the effects of *GmRALF22* on the growth of soybean under LP condition, a vector containing *GmRALF22* (pBEs-GmRALF22) was constructed and transferred into *B. subtilis* RIK1285 as described (Tang *et al*., 2022). Protein hybridization test showed that the transformed *B.subtilis* could secrete GmRALF22 into media (Fig.S4). Under LP conditions, soybean roots were inoculated with either *B. subtilis* that contains empty vector or *B. subtilis* that harbors *GmRALF22 (B. subtilis*-GmRALF22), respectively. Our results showed that inoculation of *B. subtilis*-GmRALF22 improves the vegetative growth of soybean in LP (Fig. 6A). On the 14st and 21st day after inoculation of *B. subtilis*-GmRALF22, in contrast to the inoculation of *B. subtili*s-empty vector, the height of soybean is increased by 12.27% and 24.26% respectively after inoculation with *B. subtili*s-GmRALF22 (P < 0.05, Fig. 6B). On the 21st day after inoculation with *B. subtili*s*-*GmRALF22, the fresh weight and dry weight of soybean in LP are significantly increased by 80.76% (Fig. 6C) and 55.41% (Fig. 6D), respectively. To investigate the effects of *B. subtilis* inoculation on soybean under LP stress, we carried out a treatment with *B. subtilis*-empty vector and no inoculation of *B. subtilis* under both HP and LP conditions. As shown, no significant differences were observed in terms of height, FW and DW of soybean between uninoculated and inoculated of *B. subtilis*- empty vector (Fig. 6). This suggests that the secretion of GmRALF22 protein by *B. subtilis* plays positive roles in alleviating soybean stress under LP conditions.

**Fig. 6.**
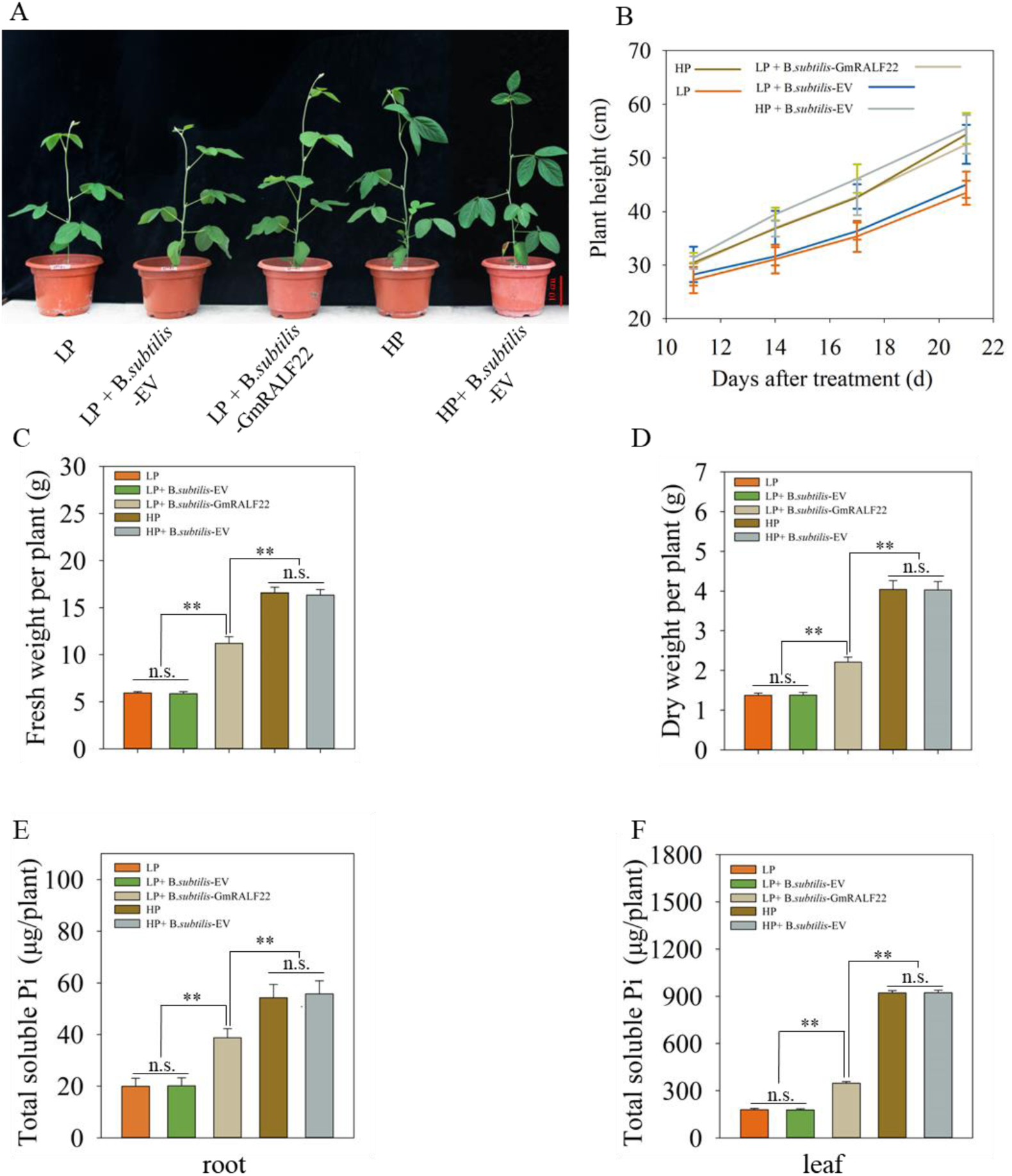
Bacterial colonization and delivery of GmRALF22 by *B. subtilis* alleviates LP stress. (A) The growth performance of soybean colonized with *B. subtilis,* which secretes GmRALF22 or not (EV, empty vector) under high P (HP) and low P (LP) conditions, respectively. (B) The plant height was measured at indicated time point after colonization of *B. subtilis,* which secretes GmRALF22, in HP and LP conditions, respectively. (C) fresh weight (FW) of the whole soybean plants;(D) dry weight (DW) of the whole soybean plant; (E-F) total soluble Pi roots and leaves. HP refers to high P conditions with a nutrient solution containing 1mM of phosphate, while LP indicates low conditions with a nutrient solution containing only 5 μM of phosphate. The FW, DW and soluble phosphate content of soybean was sampled and determined 21 days after the first inoculation of *B. subtilis*. Data are means ± SE (n = 4). The asterisks indicate significant differences; n.s., not significant (P < 0.05, Student’s *t*-test).

To verify whether GmRALF22 affects the absorption of soluble Pi (SPi) in soybean, we measured and calculated the total SPi of soybean under different treatments. In LP, compared with inoculation with *B. subtilis*-empty vector, inoculation of *Bacillus subtilis* GmRALF22 effectively increases the contents of SPi in soybean roots and leaves, and the total SPi in soybean leaves and roots increased by 85.47% and 142.91% respectively, (Fig. 6E, F). Similarly, in HP and LP, there was no significant difference in the total amount of solubility between uninoculated and inoculated *B. subtilis*-empty vectors. Thus, we argue that GmRALF22 appears to enhance soybean’s tolerance to LP by facilitating Pi absorption.

### 3.6 Inoculation of B. subtilis-GmRALF22 up-regulates of the expression of GmPTs

Inoculation of *B. subtilis*-GmRALF22 increased the content of SPi in soybean roots and leaves (Fig. 6E, F). To further elucidate the underlying molecular mechanisms, we used qRT-PCR to detect the relative expression levels of *Glycine max* Pi transporter genes (*GmPTs*) in soybean roots inoculated with *B. subtilis*-empty vector and *B. subtilis*-GmRALF22. The results showed that inoculation of *B. subtilis*-GmRALF22 increases the relative expression levels of *GmPT2, GmPT7* and *GmPT11* by 0.70, 0.85, 2.27 times under HP conditions and 0.71, 5.65, 2.07 times under LP compared with *B. subtilis*-empty vector (Fig. 7A, B, C). However, the expression of *GmPT13* and *GmPT14* was significantly up-regulated when inoculated *B. subtilis*-GmRALF22 under HP condition, and increased by 4.93 and 41.3 times respectively, but there was no significant change under LP condition (Fig. 7D, E).

**Fig. 7.**
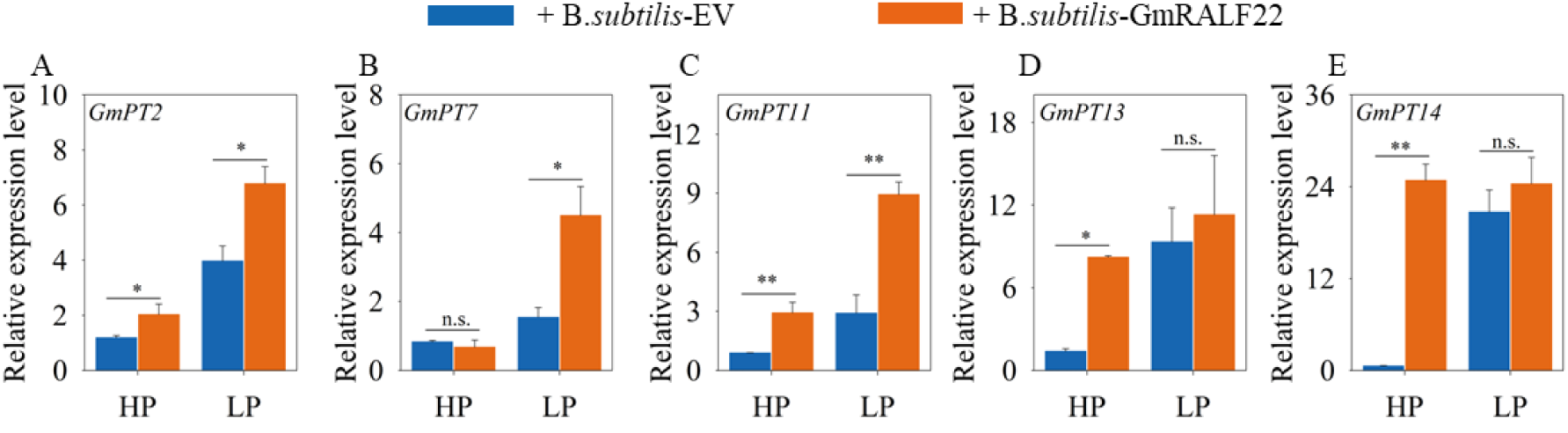
*B.subtilis* colonization and delivery of GmRALF22 increases the expression of soyeban high affinity phosphate transporters genes (GmPT) in roots. (A-F) *B. subtilis* with GmRALF22 and empty carrier were inoculated in HP and LP soil, respectively. After 21 days, RNA was extracted from soybean roots to detect GmPT transcription. The housekeeping gene *GmEF1a* (*Glyma.17G186600*) was used to normalize the qRT-PCR results. HP, 1mM phosphate; LP, 5 μM phosphate. Data are means ± SE (n = 4). The asterisks indicate significant differences; n.s., not significant (*, P < 0.05; **, P < 0.01; Student’s *t*-test).

### 3.7 GmRALF22 enhances Arabidopsis primary root growth in LP

To explore the effects of GmRALF22 on plant root growth, we synthesize mature GmRALF22 protein via Sangon Biotech (China, www.sangon.com). In this study, wild-type *Arabidopsis thaliana* (Col-0) and *Atfer4* mutants (T-DNA insertion material) cultured in normal 1/2 MS medium for 5 days in HP and LP. Synthetic GmRALF22 was applied to the Col-0 and *fer4* roots, respectively (Fig. 8A). As documented, being a cell membrane localized protein kinase, *Arabidopsis thaliana* FERONIA (FER) is the receptor of RALF. FER plays important roles in *Arabidopsis thaliana* growth, development, and immunity (Song *et al*.,2022; Tang *et al*.,2022; Zhu *et al*.,2020). Under HP conditions, application of GmRALF22 significantly inhibits the growth of Col-0 primary roots by 10.0% (Fig. 8B). Our results are consistent with the effects of AtRALF1 on Arabidopsis roots growth (Bergonci.,2014). Under LP, the application of GmRALF22 alleviates the inhibition of LP on the growth of Col-0 primary roots. Compared with the control, the application of GmRALF22 increases the length of primary roots by 8.73% (Fig. 8B), while the number of lateral roots and the density of lateral roots did not change significantly under either LP or HP conditions (Fig. 8C). Previous studies have shown that RALF modulates plant growth and development through FER kinase (Song et al., 2022; Tang et al., 2022; Zhu et al., 2020). In order to investigate the dependence of GmRALF’s impact on Arabidopsis primary root growth on FER, we then employed the *Atfer4* mutant. In HP and LP, the application of GmRALF22 to the roots of *Atfer4* mutant has no significant effects on the primary root length and the density of lateral root (Fig. 8B-D). This suggests that the roles of GmRALF22 in Arabidopsis primary root and lateral root growth and development is dependent on FER.

**Fig. 8.**
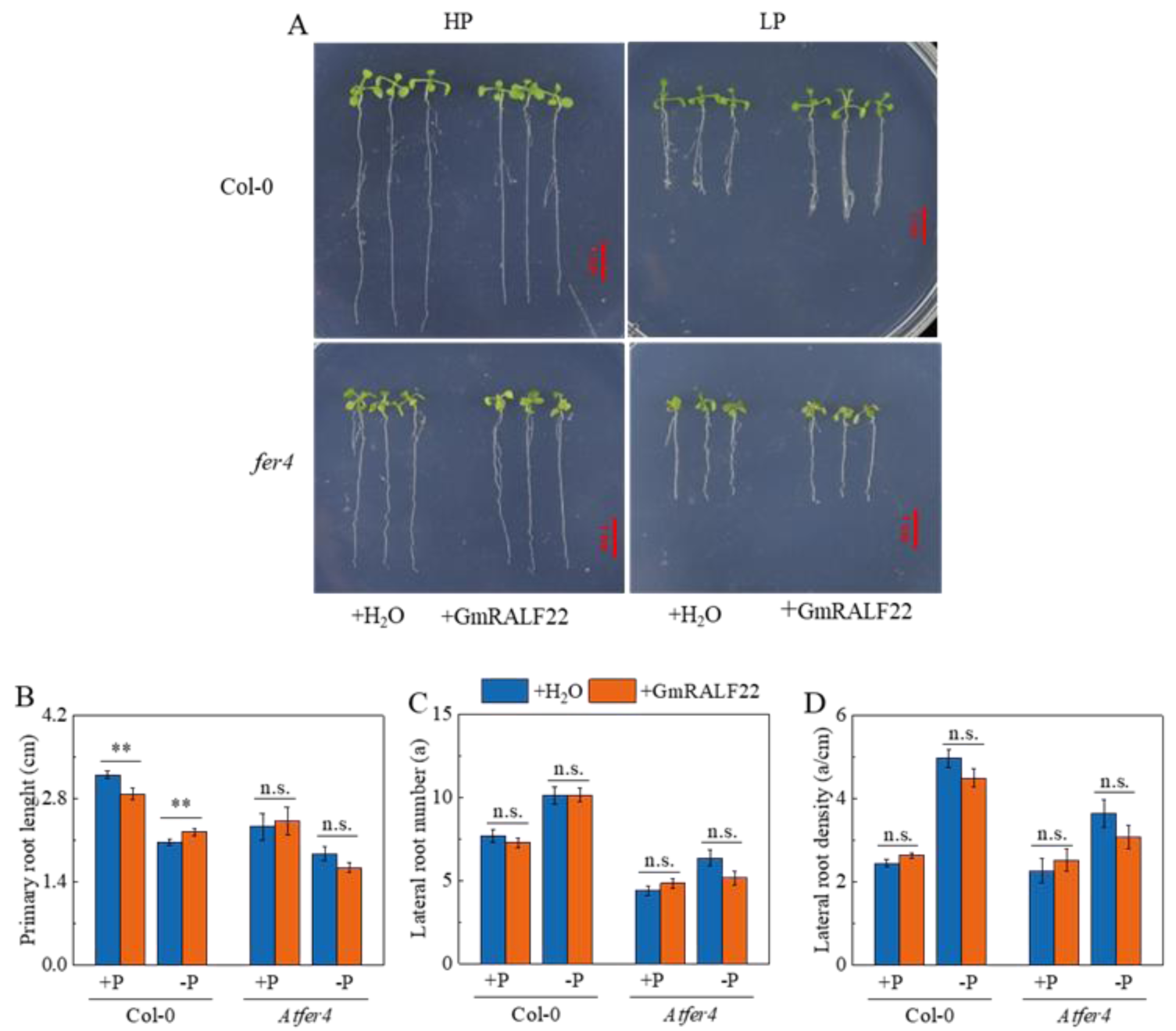
The effects of mature GmRALF22 protein on the root growth of *Arabidopsis thaliana* under high and low P conditions. (A) The effects of adding H2O and GmRALF22 protein on the root growth of *Arabidopsis thaliana* was assessed in wild-type Col-0 and *fer* mutant plants under high and low P culture conditions. Scale bar, 1cm. (B) Length of primary root; (C) number of lateral roots; (D) density of lateral roots. HP, 625 μM KH2PO4; LP, 5 μM KH2PO4. The concentration of GmRALF22 is 2 μM. Data are means ± SE (n = 4). The asterisks indicate significant differences from the HP (P < 0.05, Student’s *t*-test).

### 3.8 Transcriptomic sequencing analysis of soybean roots inoculated with B. subtilis- GmRALF22

Inoculation with *B. subtilis*-GmRALF22 was found to enhance soybean growth under LP and up-regulate the expression of Pi transporter genes (Fig. 6; Fig. 7). To further investigate the underlying molecular mechanisms, transcriptome sequencing was performed on the roots inoculated with either empty vector or GmRALF22 under LP conditions. The differentially expressed genes (DEGs) was identified based on that a false discovery rate (FDR) is less than 0.05 and a log2 fold change is greater than 1. A total of 3294 DEGs were identified, including 2041 upregulated DEGs (Table S7) and 1253 downregulated DEGs (Table S8; Fig. S2).

The differential gene expression (DEG) was subjected to Gene Ontology (GO) analysis, which is based on three aspects: biological process, cellular component, and molecular function (Fig. S3). As depicted in the figure, among the 25 classifications of biological processes, DEGs were predominantly enriched in metabolic process, cellular process, and single-organism process (GO:0044699) (Fig. S3). In terms of cell components classification, DEGs were mainly enriched in cell (GO:0005623), cell part (GO:0044464), and membrane (GO:0022804). Regarding molecular functional classification, DEGs were primarily enriched in catalytic activity (GO:0003824) and binding function (GO:0005488) (Fig. S3). A total of 888 DEGs were found to be enriched in the KEGG metabolic pathway, with a significant enrichment observed in metabolic pathways and biosynthesis of secondary metabolites (Fig. 9).

**Fig. 9.**
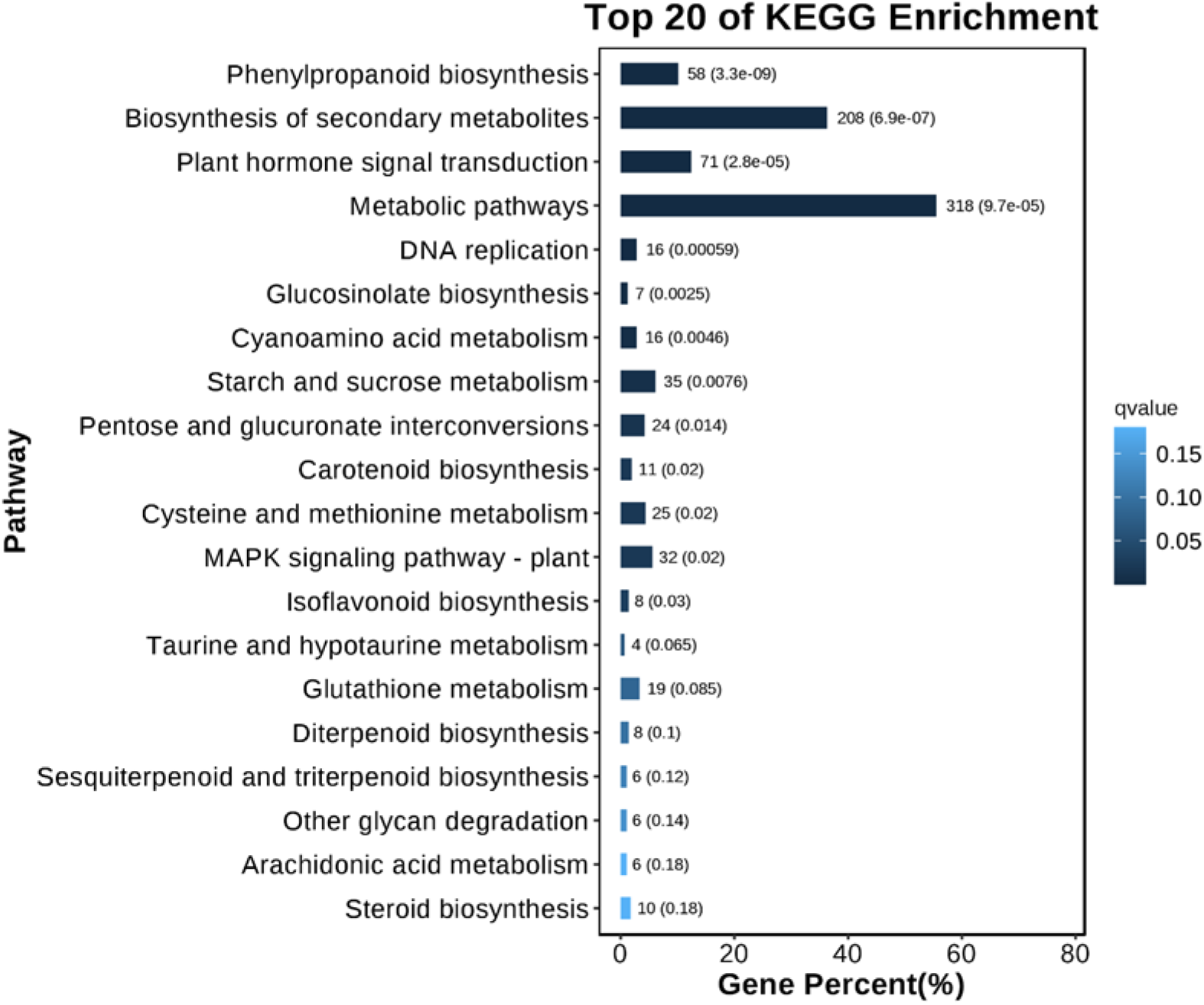
KEGG annotation of DEGs in root of *B. subtilis*-GmRALF22 vs. *B. subtilis*-EV. *B. subtilis* inoculated with GmRALF22 and empty carrier were grown in HP and LP soil, respectively. After 21 days, RNA was extracted from soybean roots for transcriptomic analysis using RNA-seq. HP: 1 mM phosphate, LP: 5 μM phosphate. Experiments were repeat three times.

We noticed that many P nutrition-related genes are found to be the DEGs many of which encode phosphatase (Table S7; Table S8). Table S3 lists the first 20 P-related and significantly up-regulated genes, among which phosphatase and phosphate synthase-related genes such as *FBPban1 (Fructose-1,6-bisphosphatase), GGPPS (Geranylgeranyl Pyrophosphate Synthase), TPS (Trehalose-phosphate Synthase), PP2C (Probable Protein Phosphatase 2C) and GPT2 (Glucose-6-phosphate/Phosphate Translocator 2)*.

In addition, it was observed that the induction of GmRALF22 resulted in up-regulation of *ITPK* (Inositol Phosphate Kinase) (Table S3). This finding suggests that the application of GmRALF22 under LP conditions not only enhances soybean’s P absorption but also promotes the expression of phosphatase genes, thereby facilitating substrate hydrolysis by phosphatases and improving soybean’s P nutrition under LP conditions. However, whether it can stimulate acid phosphatase secretion in soybean roots remains unclear.

It has been demonstrated that P nutrition is intricately linked to plant immunity (Chan et al., 2021). Strong plant immunity is observed under sufficient P conditions, whereas a decrease in plant immunity occurs in LP (Tang et al., 2022). As reported, Arabidopsis RALF modulates P nutrition (Tang et al., 2022) by regulating the immune response of plants.

Furthermore, JA and SA are revealed to regulate plant immunity (Cox, 2022; Kusajima et al., 2022). Analysis of DEG showed that a total of 12 DEGs involved in JA signal are up- regulated, ten out of which are TIFY transcription factors. As documented, TIFY are important transcription factors in JA signaling pathway. Moreover, JAR (Jasmonate Resistant) and JMT (Jasmonic Acid Carboxyl Methyltransferase) genes related to JA biosynthesis were also found (Table S4). These genes play an important role in plant immune response of JA (Shikha et al., 2023).

Interestingly, we identified 8 DEGs that are involved in SA signaling, among them, six DEGs are significantly up-regulated and two are significantly down-regulated (Table S5) by inoculation of *B. subtilis-*GmRALF22. Notably, five out of them are salicylate methyltransferases (SAMTs), which convert SA into methyl salicylate (MeSA), while the remaining three DEGs encodes salicylate binding proteins 2 (*SABP2*), which convert MeSA back to SA. As documented, both *SAMTs* and *SABP2* play crucial roles in plant SA pathway and contribute to plant resistance against adverse environmental conditions (Xu et al., 2006; Tripathi et al., 2010). Table S6 lists other immune-related genes. For example, Mycoplasma-Like Organisms (*MLO*) is related to plant disease sensitivity (Rispail et al.,2016); Ethylene-Responsive Transcription Factor (*ERF*) is an ethylene responsive transcription factor. Many studies have verified that *ERF* genes are involved in responding to various biotic and abiotic stresses in different plant species (Zhang et al., 2022). *SHN*, a branch of *ERF*, participates in plant resistance to adverse environment and pathogens by regulating plant epidermis wax and defensins (Khoudi et al.,2022). MYB is a large family of transcription factors, and many different MYB members are involved in the responses to a variety of biotic and abiotic stresses, such as *AtMYB73* positively regulates *Arabidopsis thaliana* resistance to flax leaf spot (Jia et al.,2011).

Taken together, these results suggest that GmRALF22 modulates the trade-off of plant growth and immunity via affecting JA-, SA-, or other immune-related genes, thus changing the composition of rhizosphere microbial community.

Next, ten DEGs were randomly selected for qRT-PCR analysis, including transcription factor BHLH14-like *Glyma.16G020500* and *Glyma.07G051500*; WRKY75-like *Glyma.08G011300*; JA related *Glyma.01G204400, Glyma.17G047700*, *Glyma.15G179600*, and *Glyma.07G041400*; hormone-related *Glyma.01G153300*; heavy metal transporter PDR- like ABC transporter-related *Glyma.03G168000*; glucose hexaphosphate-related *Glyma.06G032451*; and phosphatase-related *Glyma.20G000800*. When treated with *B. subtilis*-GmRALF22, *Glyma.01G204400*, *Glyma.16G020500*, *Glyma.17G047700*, *Glyma.01G153300*, *Glyma.03G168000*, *Glyma.07G051500*, *Glyma.15G179600*, *Glyma.07G041400* and *Glyma.08G011300* were up-regulated by 5.38, 6.95, 3.74, 9.00, 16.65, 59.52, 1.41,1.35 and 0.23 times, respectively, while the transcripts of *Glyma.06G032451* and *Glyma.20G000800* are decreased by 57.61% and 72.44% in contrast to the control (no GmRALF22 application) (Fig. 10).

**Fig. 10.**
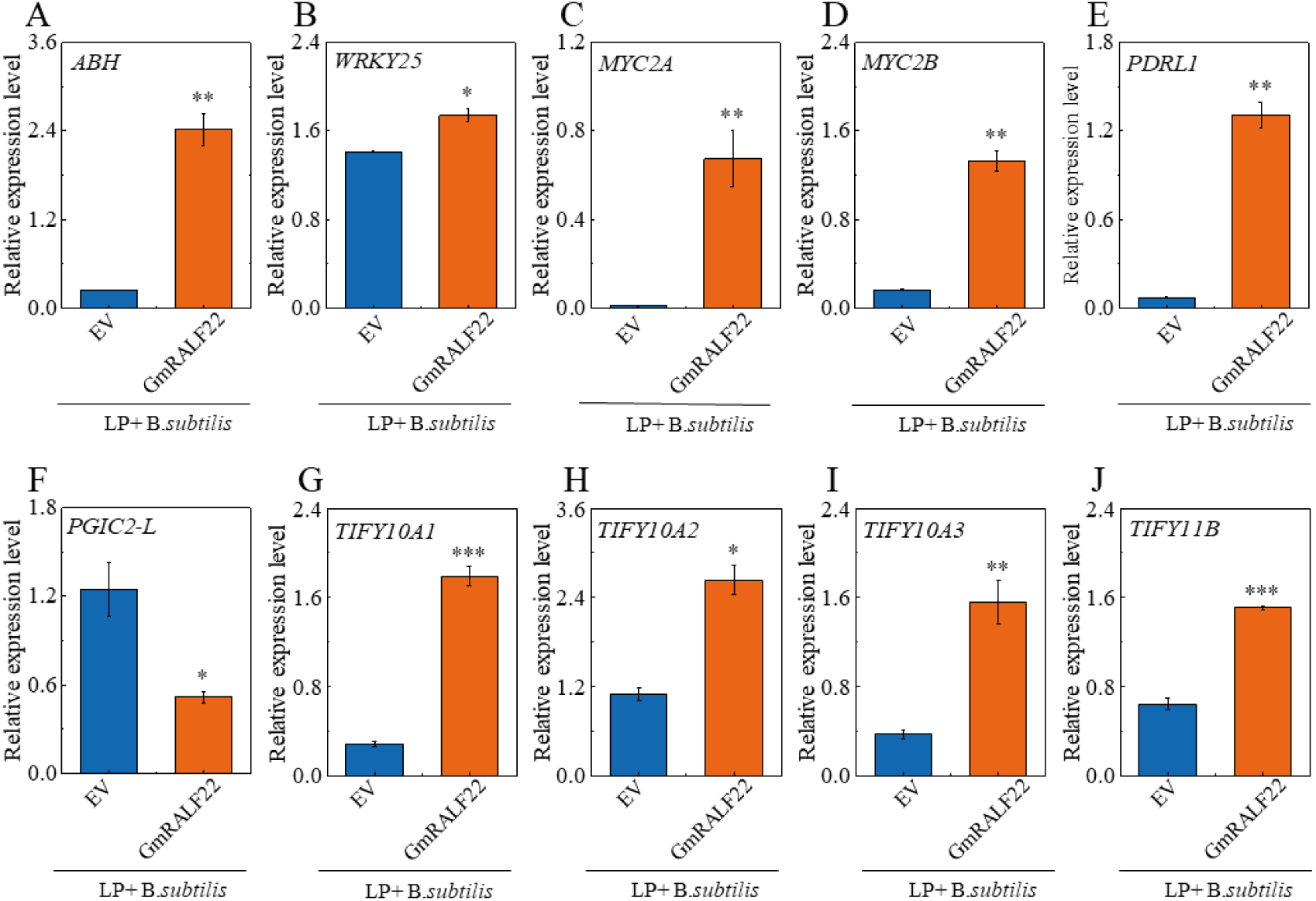
The expression profiles of some DEGs in soybean roots revealed by qRT-PCR. (A-D) The expression levels of four transcription factor genes, including one *ABI* (*ABH: Glyma.01G153300*), one *WRKY* (*WRKY25: Glyma.08G011300*), and two *MYB* genes were comparatively analyzed (*MYC2A: Glyma.07G051500*; *MYC2B*: *Glyma.16G020500*). (E-F) Two genes related to root development (*PDRL1: Glyma.03G168000*; *PGIC2-L*:*Glyma.06G032451*). (G-J) Four genes related to JA signal response (*TIFY10A1*: *Glyma.01G204400*; *TIFY10A2*: *Glyma.15G179600*; *TIFY10A3*: *Glyma.07G041400*;*TIFY11B*: *Glyma.07G041400*). The reference gene is *GmEF1a* (*Glyma.17G186600*). Data are means ± SE (n = 4). The asterisks indicate significant differences; n.s., not significant (*, P < 0.05, **, P < 0.01; ***, P < 0.001, Student’s *t*-test).

## 4. Discussion

Plants have evolved sophisticated mechanisms to cope with the everchanging environment. As reported, small molecular peptides that are sensed by cell membrane- localized receptor like kinase play crucial roles in regulating plant growth, development, and stress responses (Ge et al., 2019; Zhao et al., 2021; Tang et al., 2022). These peptides act as short- or long-distance mobile signal molecules that establish an intercellular signaling network (Liao et al., 2017; Murphy et al., 2012).

RALF is one kind of peptides that is abundant in cysteine. RALFs have been demonstrated to induce the alkalization of the surrounding environment within tobacco cell suspension culture medium, leading to an elevation in pH (Pearce et al., 2001). *RALF* genes have been extensively studied in model plants such as Arabidopsis, tobacco, tomato, and moss. Up to date, RALFs have been revealed to modulate root development and morphogenesis (Ginanjar et al., 2022; Wu et al., 2007; Zhu et al., 2020), sexual reproduction (Covey et al., 2010), acidification (Yang et al., 2022), cell expansion (Zhang et al., 2020), and plant immune response (Tang et al., 2022). Soybean is important crop for providing human being with oil and protein. However, the functions of *RALF* genes in soybean remains poorly understood.

In this study, bioinformatics analysis showed that soybean genome has 27 *RALF* genes (Table S2). Evolutionary analysis demonstrated that GmRALFs could be classified into three subgroups (Fig.1), with *GmRALF22* belonging to subgroup III and exhibiting high homology with Arabidopsis AtRALF24 and AtRALF31 (Fig.1). However, no studies on *AtRALF24* and *AtRALF31* have been reported. Moreover, three conserved motifs are found in the soybean RALFs, and GmRALF22 contains all three motifs (Fig.S1). *GmRALF* genes have tissue- specific expression patterns, indicating their crucial role in root, nodule, flower, and seed development (Fig. 3A). Hence, functional studies on the *RALF* genes that are highly expressed in soybean nodules can shed fresh light on their role in regulating nodulation and nitrogen nutrition (Collier et al., 2012).

Based on previous RNA-seq studies, eleven soybean *RALF* genes (*GmRALF1, GmRALF5, GmRALF6, GmRALF9, GmRALF11, GmRALF14, GmRALF21, GmRALF22, GmRALF25, GmRALF27)* respond to LP stress in roots and leaves (Fig. 3B) (Mo et al, 2019; Zeng et al, 2018). Here, we verified that the expression levels of *GmRALF10, GmRALF11*, and *GmRALF22* were up-regulated by LP in both roots and leaves (Fig.4D, E, G), while *GmRALF5, GmRALF6*, and *GmRALF25* were significantly upregulated by LP in leaves (Fig.4B, C, H). We also noticed that some Arabidopsis RALF genes are induced by LP (Tang *et al*., 2022)

In P-deficient conditions, the MYB domain transcription factor *PHR1* is constitutively expressed, and directly induces the expression of *Phosphate Transporter 1* (*PHT1)* to promote Pi uptake (Yang et al., 2024). Apart from its role in controlling Pi transport, PHR1 also inhibits plant defense by activating JA biosynthesis (Castrillo et al., 2017). The PHR1 protein binds to specific elements (P1BS, GNATATNC) of Pi starvation-inducible genes including PHT1, phosphatase and RNase genes (Sun et al., 2015). As reported, Arabidopsis PHR1 binds the P1BS element in the promoter region of *AtRALF*s (Tang et al., 2022). In line with this, we discovered that the promoter region (-2000 bp) of *GmRALF5, GmRALF6, GmRALF10, GmRALF11, GmRALF20, GMRALF22, and GmRALF25* contain the P1BS *cis*- element (Fig.4A). GmPHR1 has been found to be P signaling-related transcription factor. Dual-luciferase assay and EMSA analysis demonstrated that GmPHR1 directly binds to the P1BS element of *GmRALF22* promoter and stimulates its expression (Fig. 5). In Arabidopsis *AtRALF* genes such as *AtRALF4, AtRALF22*, *AtRALF33* and *AtRALF34* containing P1BS elements are responsive to LP conditions (Tang et al., 2022), indicating a conserved mechanism for plant *RALFs* responding to LP through the PHR1 pathway. However, whether other soybean *RALFs* respond to LP stress remains unclear. This requires further investigation. Additionally, there are other PHR-like transcription factors (PHLs) present in the soybean genome. Thus, it is worth exploring whether other PHL directly regulates the transcription of *GmRAL22*. In future, creating soybean loss of function of *PHR1* mutants to detect the expression levels of soybean *RALF* in LP will provide genetic evidences.

Previous studies have revealed that *RALFs* play crucial roles in regulating plant root growth and development (Zhu et al., 2020). For instance, Arabidopsis’s RALF promotes root hair growth by facilitating FER-mediated eIF4E1 phosphorylation along with targeting modules involving ROP2 and RS14 (Zhu et al., 2020). Additionally, AtRALF34 influences lateral root development through its participation in cell division within the middle column sheath (Murphy et al., 2016). Furthermore, interaction between FER phosphorylation RIPK inhibits root growth in Arabidopsis via modulation by RALF1 (Du et al., 2016).

Although RALF protein has a small number of amino acids, several critical motifs are essential for its function, for example, the motif composed of two basic amino acids in RALF, such as RRIL and RRSL, which serves as the cleavage site of AtS1P protease (Matos et al., 2008). The N-terminal YISY motif is crucial for binding to RALF receptors and alkalinization (Xiao et al., 2019). Four conserved amino acids and RCRR motifs at C-terminal play an important role in the activity of RALF peptides (Abarca et al., 2021). In this study, mature GmRALF22 protein (excluding amino acids before S1P protease cleavage site) was synthesized to investigate its effect on root growth. Under LP condition, application of mature GmRALF22 significantly increased primary root length in Arabidopsis Col-0 but not in mutant *Atfer4.* (Fig. 8), suggesting that GmRALF22 regulates primary root growth under LP condition via interacting with FER receptor kinase at least in part. However, the mechanism underlying how GmRALF22 promotes primary root growth under LP remains unclear. Further studies are needed to determine whether GmRALF22 promotes cell division or elongation in root tip in LP.

Several studies have demonstrated that the RALF-FER pathway primarily responds to nutrient and stress stimuli rather than defense mechanisms (Kim et al., 2021). For example, in *Arabidopsis thaliana*, the RALF1-FER pathway prioritizes high temperature response, inhibits primary root growth, responds to LP stress, and subsequently produces an immune response against adverse microbial environments (Castrillo et al., 2017; Kim et al., 2021). We uncovered that exogenous application of synthetic GmRALF22 protein impedes primary root growth under HP, but promote it in LP (Fig.8A), indicating its effects on primary root growth is dependent on P level

Under LP, soybean demonstrate increased efficiency in utilizing P, as evidenced by greater root volume, number of root tips, surface area and length (Zhou et al., 2021). AtRALF1 can bind to FER receptor kinase that locates in the cell membrane, while FER further interacts with RIPK receptor kinase to respond to RALF1 signal and inhibit primary root length in Arabidopsis (Haruta et al., 2014). There are 19 *RALF* genes in Arabidopsis, which all inhibit primary root growth (Sharma et al., 2016). Our results are consistent with previous studies. Considering GmRALF22 inhibits primary root growth under HP conditions (Fig.8A, B), thus we reason that GmRALF22 may modulate soybean root architecture via the RALF-FER signaling pathway to adapt to varying environmental P level. In HP conditions, plants can acquire sufficient nutrients from their surroundings, leading to increased biomass production as more nutrients are allocated above ground and increase immunity; conversely, in LP, plants enhance root development by increasing primary root lengths or lateral root numbers/densities while reducing overall diameter in order to expand contact areas between soils and roots, thus obtain necessary nutrients for normal growth maintenance (Gutiérrez-Alanís et al., 2018; Haling et al., 2016; Lambers et al., 2006; Niu et al., 2013; Sun et al., 2018). Moreover, the immunity of plants in LP is decreased, thus plants recruit more beneficial microorganisms to increase the nutrient bioavailability in soils (Tang et al., 2022). Here we found that GmRALF22 inhibit primary root growth under HP conditions but promote it under LP conditions (Fig.8A, B). The regulatory effect of GmRALF22 on primary root growth in Arabidopsis is dependent on the FER receptor, irrespective of HP or LP conditions (Niu et al., 2013; Sun et al., 2018). This suggests that soybean RALF may also exert a physiological role by binding to the FER receptor. Further investigations are required to explore the direct impact of synthetic GmRALF22 protein on soybean root and root hair growth and development, thus providing more solid evidences. In addition, soybean genome harbors several genes that encode the homologous of Arabidopsis FER receptors, indicating the existence of GmRALF-FER module. As documented, Glyma.18g215800 is the homolog of AtFER, and GmRALF4, GmRALF5, GmRALF24, and GmRALF25 interact with Glyma.18g215800 (Liu et al., 2022).

Recent years, more studies revealed that microorganism fertilizers are environment friendly in agriculture. In this study, we firstly confirmed the secretion of GmRALF22 protein by *B. subtilis*-RALF22 lines (Fig. S4). We then demonstrated that soybeans roots inoculated with transgenic *B. subtilis* secreting GmRALF22 exhibits a significant increase in soybean height (Fig. 6B), FW and DW in LP (Fig. 6C, D). The total SPi in soybeans is significantly increased upon application of GmRALF22-secreting transgenic *B. subtilis* (Fig. 6E, F). These findings suggest that GmRALF22 protein has potential for enhancing plant growth in LP in agriculture. Further investigation is required to elucidate the precise mechanisms through which GmRALF22 promotes soybean growth under such circumstances. Previous studies have demonstrated that RALF predominantly interacts with the FERONIA (FER) receptor kinase and primarily modulates plant immune responses (Stegmann et al., 2017; Zhang et al., 2020). In Arabidopsis, LP stress induces the expression of *AtRALF22* and elevates its protein levels. Subsequent investigations have revealed that AtRALF22 regulates the rhizosphere microbial community through the PHR1-RALF-FERONIA pathway, leading to significant enrichment of *Pseudomonas* by *B.subtilis* AtRALF22 in Arabidopsis; thus facilitating Pi uptake (Tang et al., 2022). Our study confirms that GmPHR1 binds to the P1BS element on the *GmRALF22* promoter, potentially promoting *GmRALF22* transcription (Fig. 5). Future studies are needed to determine soybean FER like protein that bind with GmRALF22.

In Arabidopsis, Flagellin Sensing 2 (FLS2) triggers a series of immune responses including activation of mitogen-associated protein kinase (MAPK) cascades, reactive oxygen species (ROS) bursts, and induction of defense response gene expression (Macho et al., 2014). The pathogen-associated molecular pattern-triggered PTI is mediated by Brassinosteroid Insensitive 1-Associated Receptor Kinase 1 (BAK1) (Dressano et al., 2017). Under LP conditions, FER acts as a receptor to perceive RALF22 and disrupts the complex formed by FLS2 and BAK1, thereby inhibiting immunity in Arabidopsis (Wu et al., 2019). These findings suggest that the GmRALF may also play a role in alleviating P deficiency through a similar mechanism. In Arabidopsis, under low nitrogen conditions, nutritional stress takes precedence over RALF1-mediated inhibition of cell growth (Fu et al., 2020; Wang et al., 2018). These findings are consistent with our results in this study, which demonstrate that GmRALF22 primarily promotes primary root growth rather than inhibiting primary root length in LP (Fig. 8B).

P transporters (PT) are crucial for Pi absorption and transportation in plants. Under LP conditions, *GmPT2* and *GmPT7* have higher expression levels in roots than other parts, while *GmPT11* is significantly induced in root, stem, and leaf but decreases sequentially in stem, leaf, and root (Qin et al., 2012). Additionally, *GmPT13* and *GmPT14* show the highest expression levels in leaves and stems under LP conditions (Qin et al., 2012). In this study, we showed that inoculation of *B. subtilis*-GmRALF22 increases the total SPi in soybean and promotes soybean seedling growth in LP (Fig.6A). Therefore, we speculated that the application of *B. subtilis*-GmRALF22 affects the expression of Pi transporter genes in soybean. To verify this, qRT-PCR analysis was carried out on soybean samples with inoculate *B. subtilis*-EV and *B. subtilis*-GmRALF22 under HP and LP conditions. Compared with *B. subtilis*-EV, the relatively expression levels of *GmPT2*, *GmPT7* and *GmPT11* are significantly induced by inoculating with *B. subtilis*-GmRALF22; and the transcripts of *GmPT2, GmPT7 and GmPT11* in LP conditions was much higher than that under HP conditions (Fig. 7A, B, C). Based on these results, we conclude that GmRALF22 regulates P nutrition via enhancing phosphate transporter expression and promoting Pi absorption and transport.

LP stress increases plant root exudates such as malate and citrate that activates different forms of P in soils, thus improving P utilization efficiency (Lyu et al., 2016; Shen et al., 2011). In addition, plants convert insoluble P into available P by increasing the activity of phosphatase under LP conditions (Li et al., 2014; Richardson et al., 2011; Wang et al., 2020). Roots can secrete protons to regulate pH in rhizosphere soils, while the increase of pH helps to dissolve calcium phosphate and other substances, reduce the P adsorbed by soil calcium, increase the content of soluble P, and then reduce the adsorption of P on the soil surface (Jones et al., 2009). We demonstrate that GmRALF22 exerts a significant positive impact on primary root development in LP environments (Fig. 8). However, whether GmRALF22 enhances soybean’s P nutrition through increasing the secretion of acid phosphatase and organic acids need to be explored.

To further investigate the underlying molecular mechanisms, we thus conducted transcriptome sequencing on the roots of soybean inoculated with *B. subtilis-*EV and *B. subtilis-*GmRALF22 under LP conditions. A total of 3294 DEGs were identified from the transcriptome data (Fig. S2). The analysis of DEGs revealed a significant up-regulation of phosphatase-related genes by inoculated with *B. subtilis-*RALF22 (Table S3). The findings imply that GmRALF22 may enhance soybean’s P nutrition by modulating the activities of phosphatase and phospholipase, thereby increasing plant-available phosphorus levels in both soybean plants and their rhizosphere.

The involvement of JA and SA in plant immune response has been reported (Cox, 2022; Kusajima et al., 2022). In Arabidopsis PHR1 interacts with JAZ (Jasmonate Zim) to regulates JA signal pathway under LP condition (He et al., 2023). The signal pathway of RALF-FER can respond to LP signal by participating in the immune response process (Tang et al., 2022). In this study, we identified twelve up-regulated JA-related DEGs (Table S3), six up-regulated and two out of 8 down-regulated SA-related genes (Table S4). In addition, many immune-related genes were found to be DEG (Table S5). Hence, GmRALF22 seems to alter Pi uptake in soybean through regulating the expression of JA, SA, or other immune-related genes. Figure 11 outlines the working model of GmPHR1-GmRALF22 in modulating P nutrition. Taken together, our results provide useful cues to decipher the function and application of soybean RALFs.

**Fig.11.**
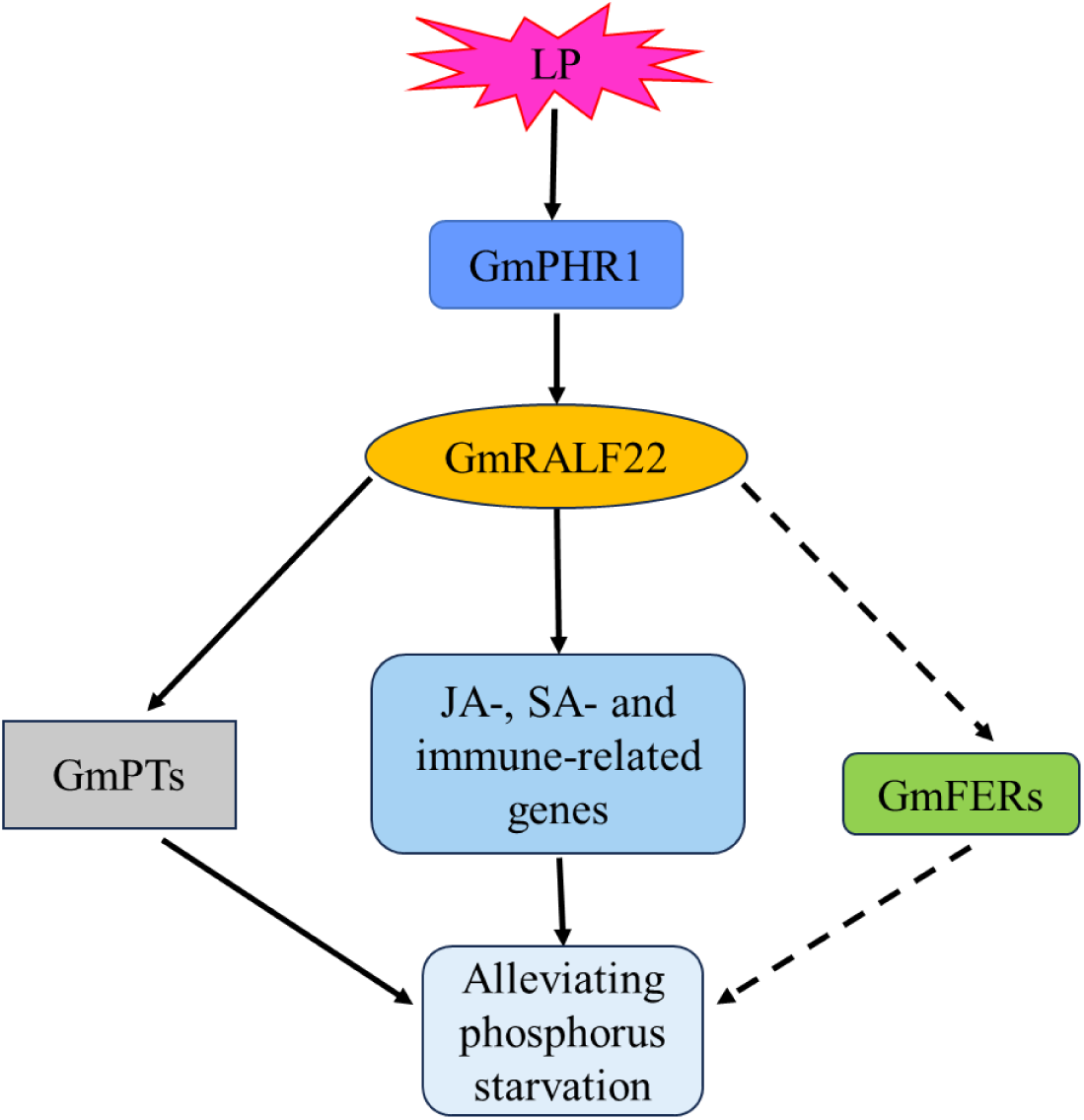
The working model of GmRALF22 in regulating phosphorus stress. The expression of the *GmRALF22* gene was positively regulated by GmPHR1 under LP stress. Subsequently, GmRALF22 facilitated the upregulation of phosphorus transporters (*GmPTs*), as well as JA-, SA-, and immune-related genes, thereby enhancing soybean’s tolerance to LP stress. In addition, GmRALF22 may regulate the tolerance of soybean to LP stress through the pathway of GmFERs.

## 5. Conclusion

Soybean genome has 27 *GmRALF*s. *GmRALF5*, *GmRALF6,* and *GmRALF10*, *GmRALF11*, *GmRALF22* and *GmRALF25* are induced in LP. GmPHR1 directly regulate the transcription of *GmRALF22* via binding the promoter P1BS *cis*-element. Secretion of GmRALF22 protein by *B. subtilis* promoted soybean growth in LP by improving soybean P nutrition through increased expression of high affinity Pi transporter gene. GmRALF22 regulates soybean Pi absorption by modulating JA-, SA- and immune-related genes expression at transcriptome level. That application of GmRALF22 promoted primary root growth in Arabidopsis in LP is dependent on FER receptor.

Supplementary data to this article can be found online at XXX.XXX.XXX.XXX

## CRediT authorship

Fangjian Li: Conceptualization, Methodology, Validation, Formal analysis,

Investigation, writing original draft, Writing review & editing. Cuishan Mai: Validation, Formal analysis, Investigation, writing original draft. Yan Liu: Validation, Formal analysis, Investigation, writing original draft. Yaru Deng: Validation. Lixia Wu: Investigation. Xinni Zheng: Validation. Huijing He: Validation. Yilin Huang: Validation. Zhenxi Luo: Validation. Jinxiang Wang: Supervision, Conceptualization, Funding acquisition, Project administration, Resources, Writing and editing.

## Declaration of Competing Interest

The authors declare that they have no known competing financial interests or personal relationships that could have appeared to influence the work reported in this paper.

## Data Availability

Data will be made available on request. RNA-sequencing data (PRJNA1104748) has been deposited in NCBI (https://www.ncbi.nlm.nih.gov/).

## Acknowledgments

This study was partially supported by Guangzhou Science and Technology Plan Project (Grant No. 202206010163) and the Science and Technology Planning Project of Guangdong Province of China Foundation (Grant No. 2021B1212040008). We thank Dr. Feng Yu (Hunan University) for providing *B. subtilis* transformation system and discussion, Dr. Dousheng Wu (Hunan University), Dr. Jing Tang (Hunan University) and Dr. Jiang Tian (South China Agricultural University) for their help.

Table S1. The list of primer pairs used in this study.

---

**Fig. S1.**
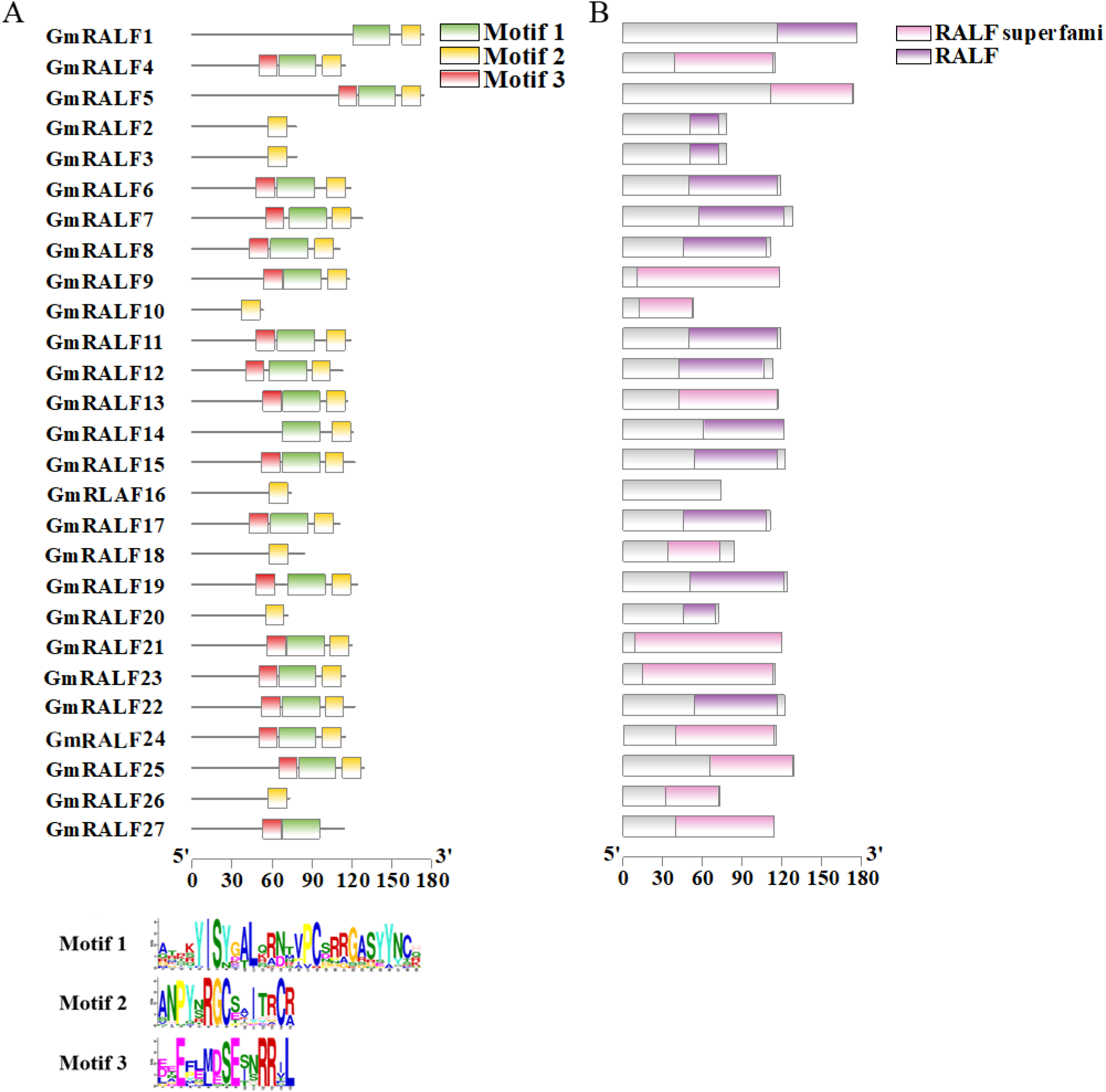
Motif analysis of GmRALF family. (A) motif; (B) protein domain. RALF superfamily/ RALF: rapid alkalizing factor family domain. (C): the amino acid sequences of conserved motifs. The letter size in the motif represents the degree of conservation.

**Fig. S2.**
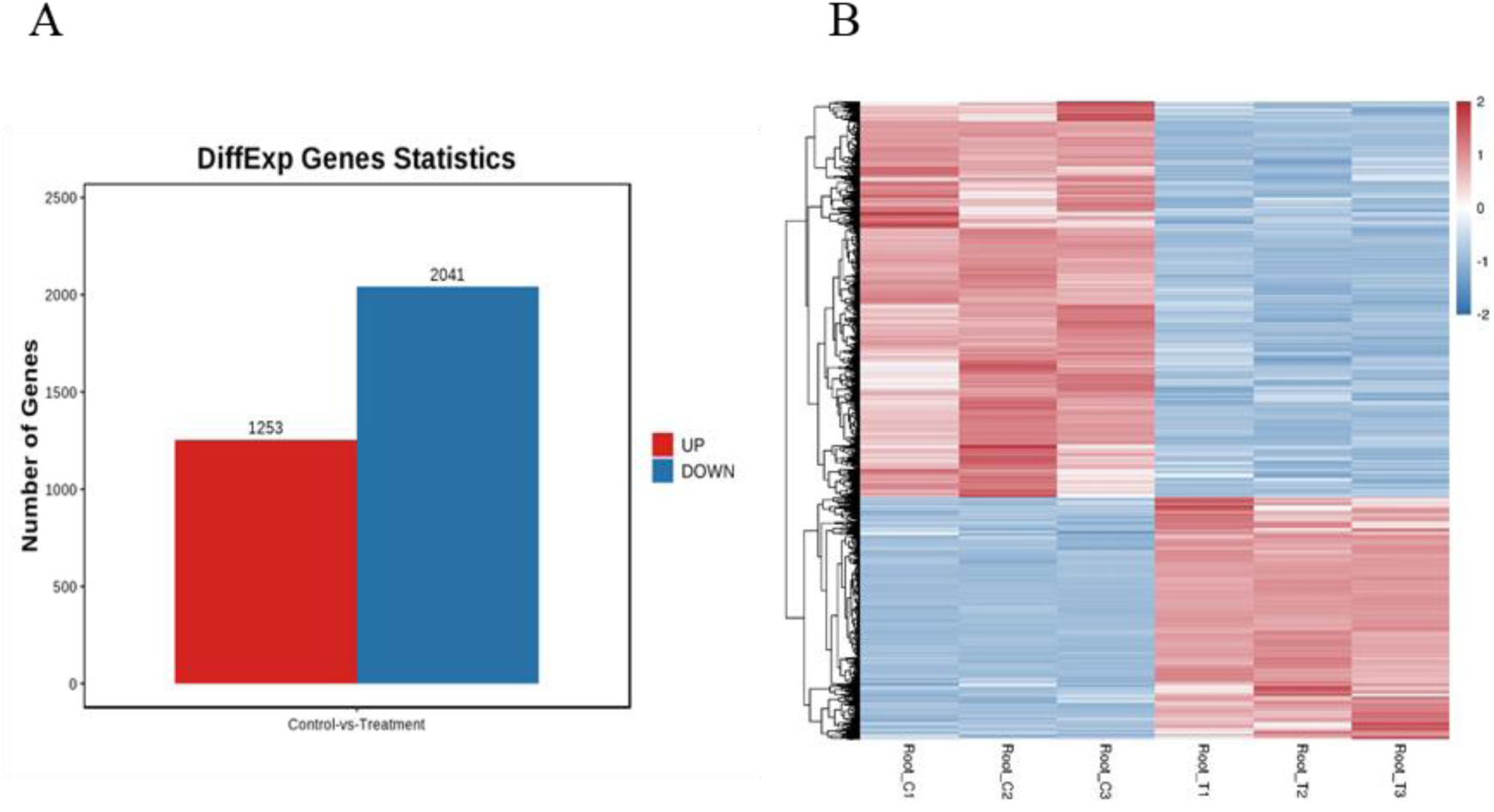
Identification of differentially expressed genes through transcriptome sequencing and visualization using a heatmap. (A) Histogram depicting the distribution of genes exhibiting up-regulation and down-regulation in transcriptome data; (B) This heatmap displays the differential expression of genes in transcriptome data, with red indicating up- regulation and blue indicating down-regulation. The intensity of color reflects the magnitude of change, with darker shades representing higher fold changes.

**Fig. S3.**
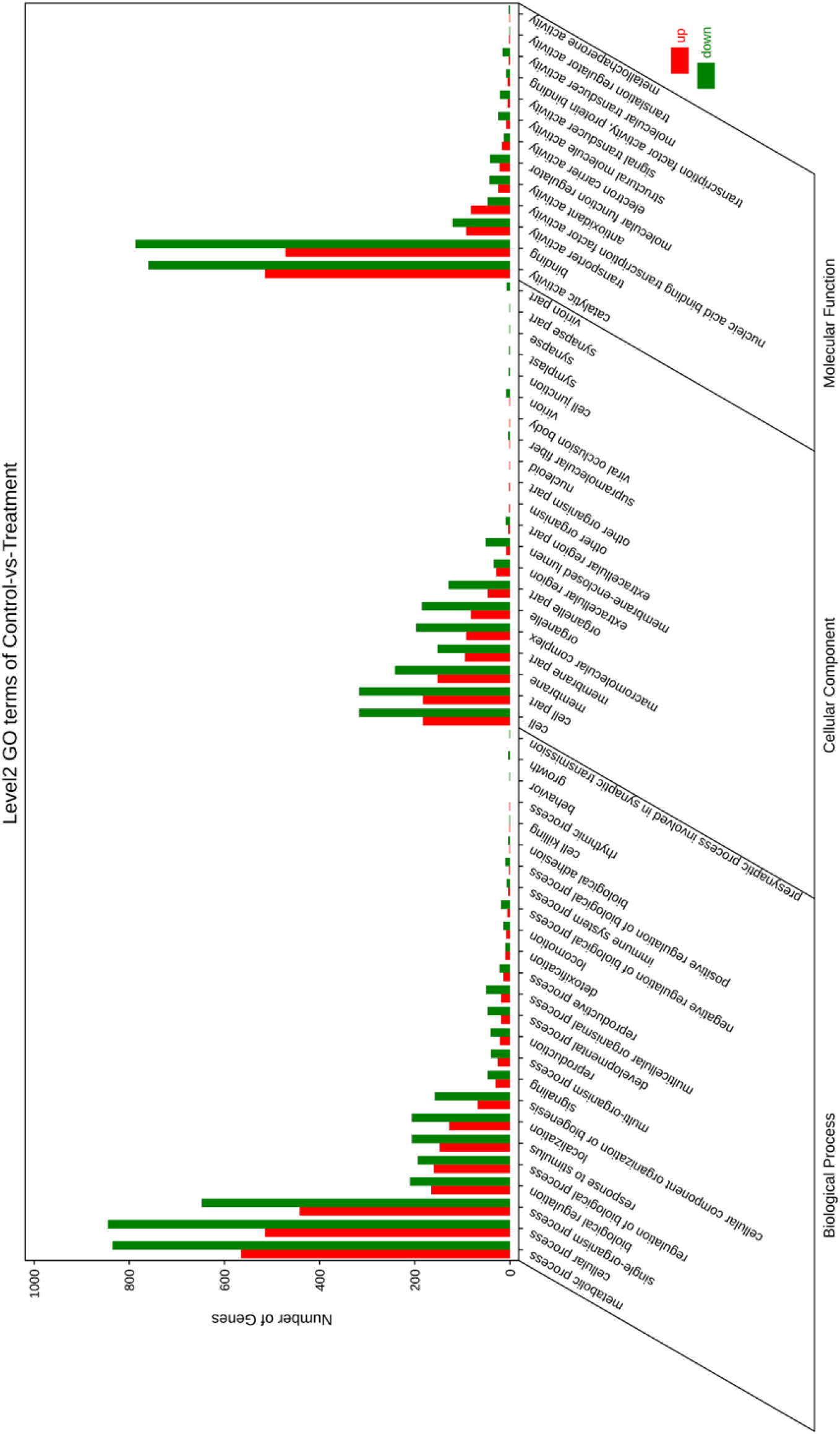
Enrichment analysis of differentially expressed genes (DEGs) was performed using Gene Ontology (GO). The abscissa represents the secondary GO classification, while the ordinate indicates the number of genes in each category. Up-regulated DEGs are shown in red, while down-regulated DEGs are shown in green. GO analysis was performed using the URL (http://systemsbiology.cau.edu.cn/agriGOv2/index.php).

**Fig. S4.**
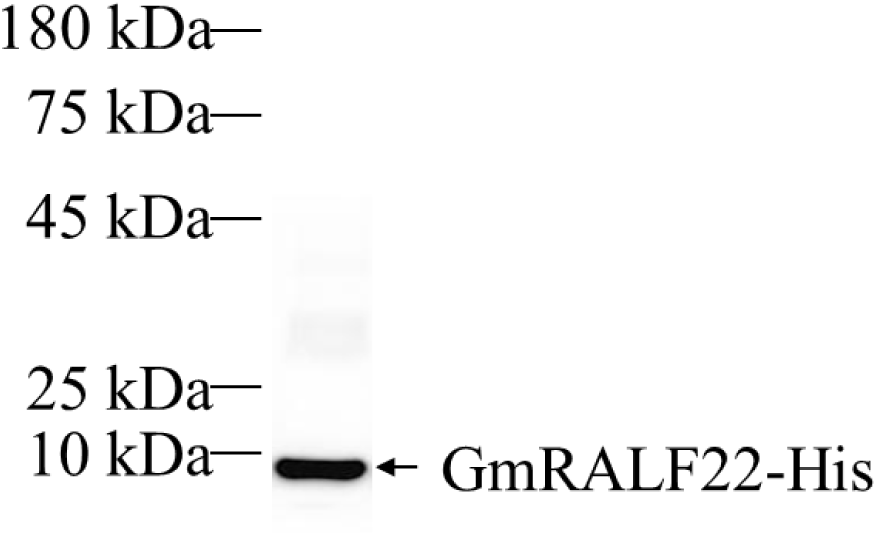
Immunoblot analysis of GmRALF22 protein in the *B. subtilis-*GmRALF22. Total protein was extracted and analyzed using anti-His antibody ((M30111, Abmart).

**Fig. S5.**
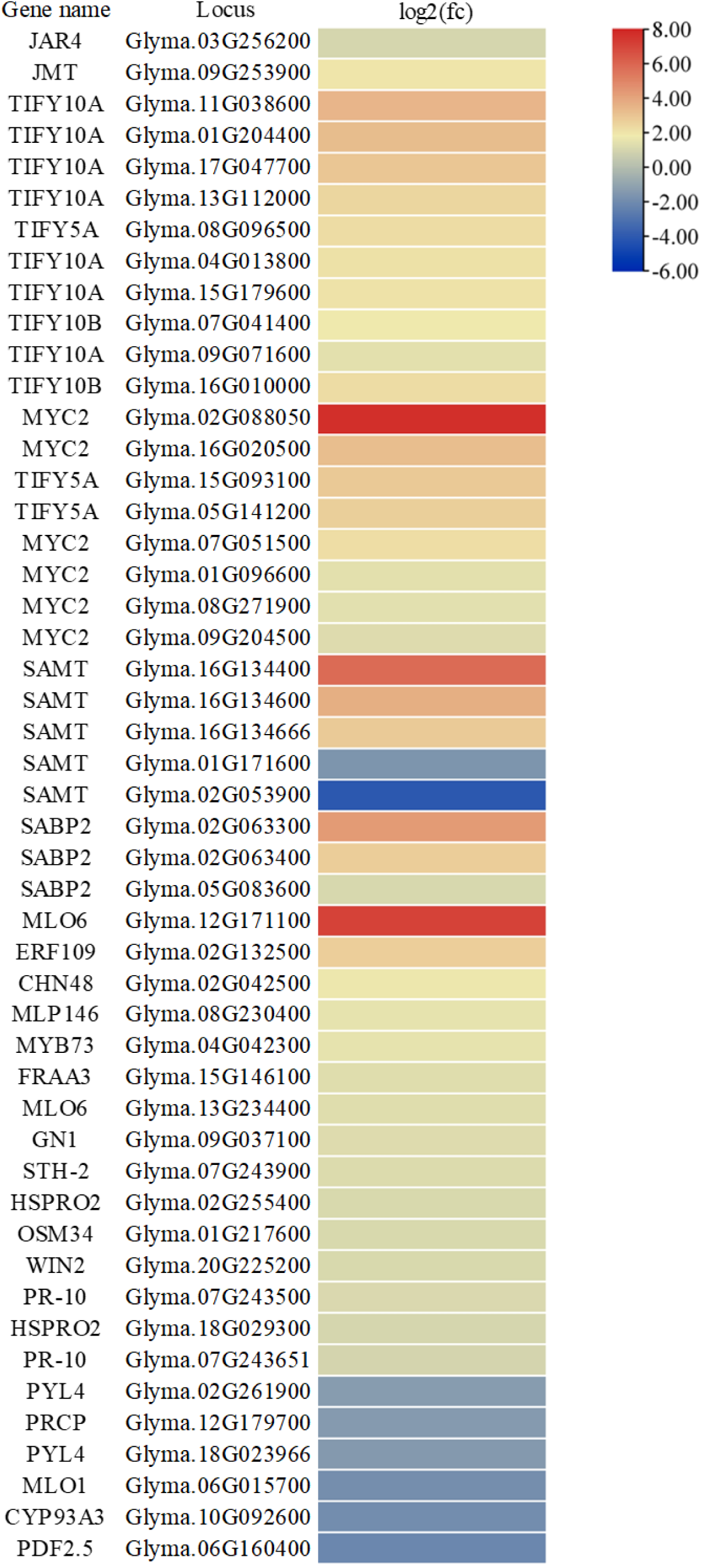
The heatmap depicts fold change of JA-, SA- and immunity-related genes that are regulated by *B. subtilis-*GmRALF22.

Table S2. General information of the GmRALF family. Note that all gene loci were obtained from the soybean database *Glycine max* Wm82.a4.v1 in Phytozome (https://phytozome-next.jgi.doe.gov/).

Table S3. P-related DEGs. Table S4. JA-related DEGs. Table S5. SA-related DEGs.

Table S6. Immune-related DEGs. Table S7. Up-regulated DEGs.

Table S8. Down-regulated DEGs.

## Notes

### Competing Interest Statement

The authors have declared no competing interest.

